# Mycobacteriophage D29-mediated lysis improves recovery of mycobacterial genomic DNA from low-biomass samples

**DOI:** 10.64898/2026.08.08.743631

**Authors:** Joseph Wambugu Gitari, Anastasia Koch, Elizabeth Kigondu, Digby F. Warner, Mandy K. Mason

## Abstract

**Background:** Detection of rare mycobacterial genotypes, including those associated with antibiotic resistance or population heterogeneity is important for diagnostic, therapeutic and research applications. This depends on efficient recovery of genomic DNA (gDNA) from sampled populations, a challenging requirement in paucibacillary clinical materials. Mycobacteria have uniquely lipid-rich, structurally robust cell envelopes which resists cell lysis by conventional methods. Here, we characterize mycobacteriophage D29-mediated lysis at the single-cell level, evaluating its utility as a biological lysis strategy for mycobacterial DNA isolation, benchmarked against the standard cetyltrimethylammonium bromide (CTAB) extraction method.

**Methods:** Conditions for mycobacteriophage D29 infection of *Mycobacterium smegmatis* (*Msm*) were established, and single-cell phage adsorption and phage-mediated lysis visualized through live-cell time-lapse fluorescence microscopy (FM). A mycobacteriophage D29-based lysis method was applied to both *Msm* and *M. tuberculosis* (*Mtb*), and extraction efficiencies compared with the standard CTAB method. Cell lysis efficiency was quantified by colony forming units (CFU), flow cytometry (FC) and FM; DNA yield was determined by quantitative polymerase chain reaction (qPCR) and droplet digital PCR (ddPCR).

**Results:** Mycobacteriophage D29 adsorption was observed at the poles and septa of individual mycobacterial cells. Phage infection was associated with loss of cytoplasmic green fluorescence protein (GFP) reporter protein, with uptake of a cell death marker propidium iodide (PI). Mycobacteriophage D29 infection resulted in a marked loss of cell viability, with >6log_10_ reduction in CFU, and cell lysis efficiencies calculated as 93.3% (FC) and 96.8% (FM). Molecular quantification (qPCR and ddPCR) indicated that the mycobacteriophage-based lysis achieved between 4- to 7-fold greater gDNA yields in *Msm* and between 3- to 12-fold greater gDNA yields in *Mtb* H37Ra compared with the CTAB method. Notably, gDNA extraction efficiencies in both mycobacterial species exceeded 92% in low-biomass samples containing approximately 100, 175 and 320 bacilli.

**Conclusion:** These results demonstrate the utility of the mycobacteriophage D29-based method for improved DNA extraction yields from mycobacteria through direct lysis of individual bacilli, with performance suited to low-biomass samples.

**Summary:** Recovering genomic DNA (gDNA) from low numbers of mycobacteria is a persistent bottleneck for diagnostics and genomic studies, because the lipid-rich mycobacterial envelope resists conventional lysis. Here we show that mycobacteriophage D29 provides an efficient, biologically selective route to mycobacterial DNA. Leveraging single-cell live imaging, we reveal that phage D29 adsorbs preferentially at the poles and septa of individual cells, and that infection is heterogeneous and asynchronous, progressing from envelope permeabilization to loss of viability. Applied as an extraction method and benchmarked against the standard cetyltrimethylammonium bromide (CTAB) protocol, phage D29-mediated lysis recovered 4- to 7-fold more gDNA in *Mycobacterium smegmatis* (*Msm*) and 3- to 12-fold more in *Mycobacterium tuberculosis* (*Mtb*). Critically, extraction efficiency exceeded 92% in both species in low-biomass samples of approximately 100, 175 and 320 bacilli, where CTAB performed poorly (<20% efficiency). These findings support phage-mediated lysis as a quantitative, near-complete DNA-recovery method that outperforms conventional extraction precisely in the paucibacillary regime of greatest clinical relevance and demonstrate the value of single-cell interrogations in building towards precision tools to engage the mycobacterial cell.

## Introduction

Bacterial populations exhibit remarkable heterogeneity, comprising various subpopulations that can be proportionally small and which may include rare, but relevant, genotypes such as those associated with antibiotic persistence, tolerance, and resistance (Jones et al., 2022). Rare genotypes are implicated in failed chemotherapy (Dewachter et al., 2019; Huemer et al., 2020; Urbaniec et al., 2022), undermining the control of many bacterial infections. Rare subpopulations such as persister cells occur at extremely low frequencies (proportions estimated between 10^-4^ to 10^-6^) in populations of bacteria (Keren et al., 2004). This is particularly relevant in the context of tuberculosis (TB) where prolonged treatment, slow and heterogeneous growth of *Mycobacterium tuberculosis* (*Mtb*), phenotypic drug tolerance, and within-host bacterial diversity may create conditions in which rare drug-refractory subpopulations can survive therapy (Coetzee et al., 2024; Xu et al., 2025). An understanding of the genetic mechanisms enabling antibiotic survivability requires detailed insight into the genotypic heterogeneity that can exist within an apparently clonal bacterial population (Boshoff et al., 2023). These investigations require techniques enabling the molecular detection and characterization of relatively small subsets of mycobacterial cells. Comprehensive detection of rare genomic variants poses a technical challenge, as this requires efficient, and unbiased gDNA extraction techniques to maximise the likelihood that chromosomal DNA from low-frequency cells within a sampled population is recovered and accessible for downstream molecular interrogation.

Conventional bacterial gDNA extraction protocols generally rely on culture expansion to generate high-biomass preparations, often comprising hundreds of millions to billions of organisms, ensuring high yields of high-quality gDNA (van Helden et al., 2001; Wilson, 2001). These bulk extraction protocols are typically optimised in terms of total DNA yield, purity and integrity rather than the proportion of individual cells successfully lysed and represented in the recovered DNA (Murase et al., 2025; Percy et al., 2024). Propagation in lab-based media is also known to impose selective pressures, altering the genetic composition of populations (Domenech & Reed, 2009; Mulholland et al., 2024), this may reduce population diversity (Metcalfe et al., 2017; Nimmo et al., 2019; Shockey et al., 2019) and alter representation of organisms due to the existence of differentially culturable bacteria (Ayrapetyan et al., 2015; Chengalroyen et al., 2016; March et al., 2025). Consequently, culture-independent extraction methods that preserve composition and diversity of the original population are an important methodological goal, particularly where clinical materials or bacillary numbers are limiting and where the requirement is to capture of rare events within a population, while minimizing or eliminating the need for *in vitro* propagation. In mycobacterial research, this is a critical consideration as samples retrieved from clinical settings frequently contain low bacillary loads (Alebouyeh et al., 2022), while there is an increasing need to detect low-frequency drug-resistant variants, or resistant subpopulations that may expand during therapy and adversely affect treatment outcomes (Cohen et al., 2012; Lozano et al., 2021). The efficient lysis and extraction of mycobacterial molecules has therefore become increasingly important with effective diagnostics increasingly dependent on accurate detection and quantification of molecules such as DNA (Chin et al., 2018; Dziri et al., 2024) and RNA (Walter et al., 2021). Inefficient lysis and release of cytoplasmic molecules from mycobacteria undermines progress in key areas including (i) the interrogation of paucibacillary mycobacterial (sub)populations via culture-free sputum, tongue swab, and aerosol sampling (Church et al., 2024; David et al., 2025; Dinkele et al., 2024; Yan et al., 2024), and (ii) the detection of ultrarare genotypes such as microheteroresistance, whose proportions can fall below the limit of detection of even advanced sequencing protocols (Chen et al., 2025; Colman et al., 2015; Colman et al., 2025).

The mycobacterial cell lysis is a known challenge in this space, with cells being relatively difficult to break open due to the complex nature of their unique cell envelope (Beaud Benyahia et al., 2024; Hashimi & Tocheva, 2024; Jacobo-Delgado et al., 2023). The mycobacterial cell envelope includes a covalently linked mycolyl–arabinogalactan– peptidoglycan complex, (Alderwick et al., 2015), a large robust macromolecular scaffold that contributes substantially to structural integrity of the cell wall (Catalão & Pimentel, 2018). A complex array of lipids, including mycolic acids makes up the outer leaflet of the outer membrane (myco-membrane), with these long-chain fatty acids aligning to form a thick, impermeable, waxy layer (Daffé, 2015), making it difficult for conventional lytic enzymes to access their molecular targets (Murase et al., 2025). Combinations of mechanical, chemical, and enzymatic lysis methods are well established in standard DNA extraction protocols, however, these techniques may not achieve optimal DNA recovery from heterogenous mycobacterial populations (Martzy et al., 2019) and may compromise DNA quality (Hall et al., 2023). The cetyltrimethylammonium bromide (CTAB) DNA isolation method, which relies on chemical and enzymatic lysis of cells, is commonly applied (Doyle, 1990; van Helden et al., 2001), however, this multi-step extraction protocol creates multiple opportunities for the loss of cells and DNA template molecules. The choice of mycobacterial lysis method therefore plays a key role in DNA isolation efficiency, which varies considerably across studies (Amaro et al., 2008; Kolia-Diafouka et al., 2018).

An alternative strategy for disrupting the structurally robust mycobacterial cell envelope is to harness lytic mechanisms that have evolved through mycobacteriophage-host interactions. Mycobacteriophage D29 (also referred to here as phage D29) is a well-characterized lytic phage with a broad mycobacterial host range (Dedrick et al., 2017; Ford et al., 1998). Following infection and intracellular replication, the phage lytic cycle culminates in disruption of the host cell and release of progeny phage together with intracellular bacterial material (Swift et al., 2020). Critically productive phage replication depends on a viable and metabolically permissive host cell, a feature that provides a means of selectively detecting viable mycobacteria (Hatfull, 2018). Classic phage amplification assays, developed as diagnostics more than two decades ago (Albert et al., 2002; McNerney et al., 2004), utilise this selective lytic activity of phage D29 to estimate viable mycobacteria in sputum, blood and milk using plaque redouts (Rees et al., 2024; Stanley et al., 2007; Swift et al., 2020). These assays were shown to be relatively rapid and effective with high sensitivity and specificity and have been developed to leverage phage D29 lytic properties for downstream molecular readouts (Beinhauerova & Slana, 2021). Tests include the commercialised Actiphage method, which eliminated the plaque-formation step by using D29 directly to lyse viable mycobacteria and, release their genetic material to be recovered from growth medium, concentrated, and purified for downstream molecular analyses (Rees et al., 2024; Swift et al., 2020). The method enabled rapid DNA recovery and PCR-based detection from bovine blood samples with low bacterial inputs (Swift et al., 2020), supporting its potential application to paucibacillary samples (Swift et al., 2020) and its compatibility with rapid, tractable and analytically sensitive molecular readouts, including loop-mediated isothermal amplification (Shield et al., 2024).

Collectively, these developments have established phage D29-mediated lysis as a versatile biotechnology for selectively accessing genetic material from viable, low abundance mycobacteria. However, despite its application in several diagnostic workflows, the adsorption, infection, permeabilization and lysis events underpinning the phage-mediated DNA release remain incompletely characterized at the single-cell level, and the relationships between bacterial input, infection efficiency, cell lysis and recoverable DNA has not been systematically defined.

In this study, we investigated phage D29-mediated lysis as an approach for DNA isolation from mycobacteria, focusing on characterizing its effects at a single-cell level and evaluating its performance across a calibrated range of bacterial cell densities. We determined the efficacy of phage infection and subsequent cell lysis in the non-pathogenic model *Mycobacterium smegmatis* (*Msm*) and compared DNA recovery with that achieved using standard CTAB extraction method which remains a widely adopted reference in mycobacterial molecular epidemiology (van Helden et al., 2001). It is selected here to provide a common benchmark against which alternative methods may be compared. Finally, we evaluated the applicability of this approach in the attenuated *Mtb* strain H37Ra.

## Results

### Phage D29 adsorbs at poles and septa and is associated with increased cell permeability

Efficient DNA recovery from paucibacillary mycobacterial samples requires lysis that is both efficient and reproducible at the level of individual bacilli. Although phage D29 has been widely used as a lytic phage, little is known about the sequence of events leading from adsorption to envelope disruption in individual mycobacterial cells. High-resolution single-cell imaging enables direct observation of phage-host interactions that are obscured in conventional population-level assays. We used this approach to determine adsorption localisation of phage D29 and how subsequent infection impacts cell envelope integrity. *Msm*::GFP cells were exposed to phage D29 pre-stained with SYTOX Orange and the infections observed by fluorescence microscopy (FM) (**Figure 1**). A total of 91 green fluorescence protein (GFP) positive bacilli were analysed. Adsorption of SYTOX Orange-labelled phage particles were observed either at one of the mycobacterial poles, both poles, or at the poles and septum (**Figure 1A**). Single pole adsorption was observed in 59% of mycobacterial cells (**Figure 1A i**), double pole adsorption in 18% (**Figure 1A ii**), and polar and septal adsorption 6% of the population (**Figure 1A iii**). SYTOX Orange fluorescence profiles confirmed the polar and septal adsorption of the phage (**Figure 1B**). No phage adsorption was observed in 17% of the GFP positive bacteria. SYTOX Orange fluorescence signal was negligible in mycobacteria not exposed to phage and in intact bacteria treated with the stain alone (**Supplementary Figure S1**). Polar and septal localisation is associated with sites of new cell wall synthesis in phages such as Fionnbharth, Muddy and Adephagia (Dulberger et al., 2023) which are the areas of active metabolism as evidenced by DMN-Tre incorporation in *Msm* cells (**Supplementary Figure S2**). These results suggest that phage D29 adsorption occurs preferentially at the metabolically active poles and septum of *Msm* bacteria and that not all cells are permissive to infection.

**Figure 1:**
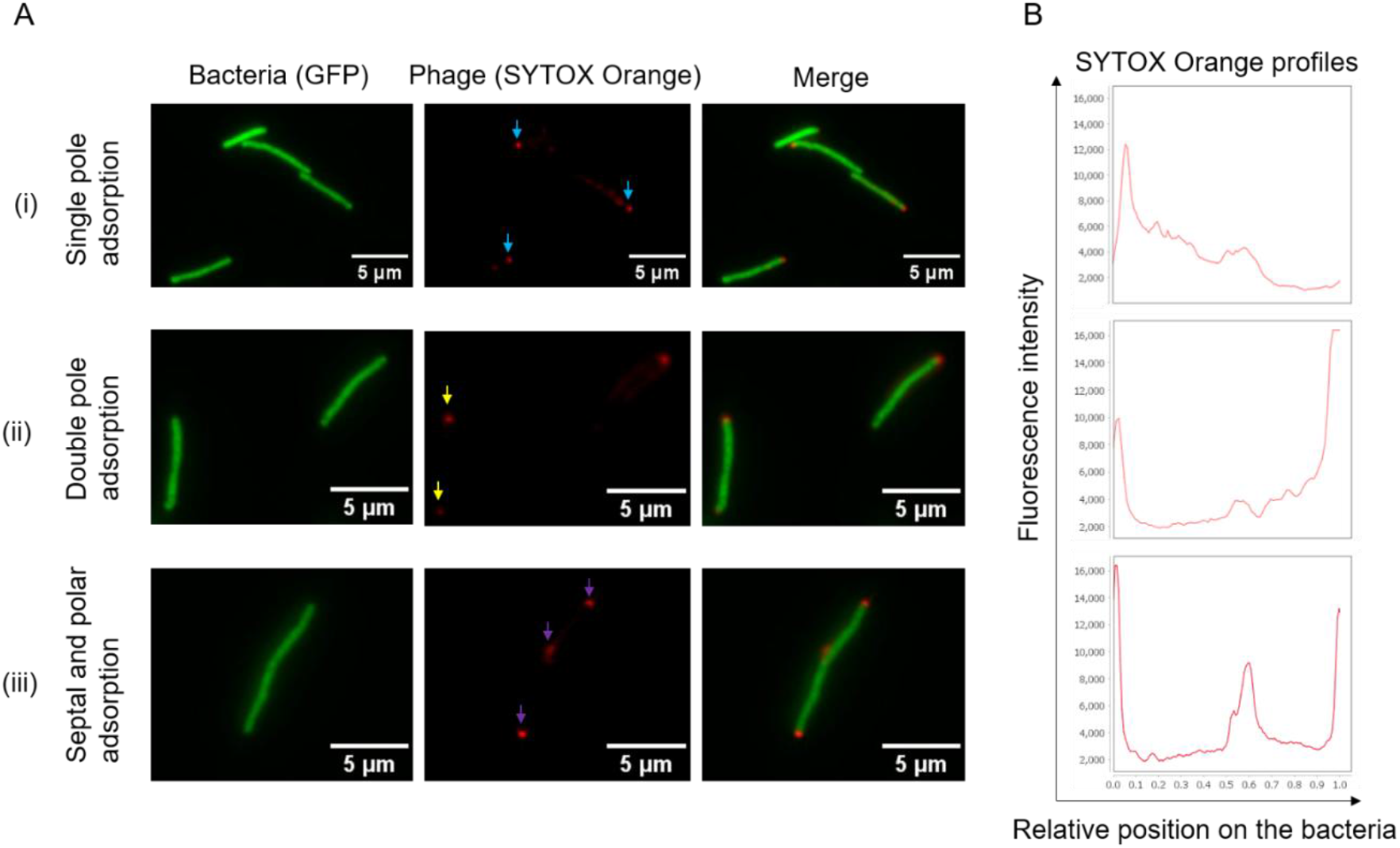
Mycobacteriophage D29 adsorbs preferentially at mycobacterial poles and septa. (A) Adsorption of SYTOX Orange-stained phage D29 on *Msm*::GFP on either single pole (blue arrows), both poles (yellow arrows) or both poles and septum (purple arrows). (B) SYTOX Orange fluorescent signal profiles mapped on standardised bacterial lengths distinguish single pole, double pole, and polar and septal signatures.

To better characterize phage-host interaction dynamics phage D29 infection kinetics were characterized at the single-cell level in *Msm*::GFP using the CellASIC ONIX2 microfluidics system. *Msm*::GFP was trapped in the microfluidic device at low density and a phage titer of ∼4.6X10^3^ plaque forming units per milliter (PFU/mL) was perfused at a pressure of 2 psi for 6 hr after which the cells were monitored for a further 17 hr under standard medium perfusion (**Supplementary Methods Figure SM1A**). Under these conditions, loss of GFP signal, a relatively stable molecule with a half-life of 26 hr (Corish & Tyler-Smith, 1999) was observed 3-9 hr into the experiment, with smooth, dark rod-shaped cells becoming grey and misshapen and absence of cell division (**Figure 2A, Supplementary video 2**). A strong GFP signal was maintained in cells that were not treated with phage over the course of the 24 hr experiment, with bacteria undergoing normal cell division (**Figure 2A, Supplementary Video 1**). The observed loss of GFP signal was supported by the phage mediated cell envelope permeabilization in the cells treated with phage where increased accessibility of BODIPY-FL-vancomycin (BODIPY-FL-van) was observed in a phage concentration dependent manner (**Supplementary Figure S3**), indicating that myco-membrane integrity is compromised with phage infection. It was notable that not all cells experienced loss of GFP at the same timepoint, possibly due to asynchronous phage infection of the bacteria. These results illustrate phage-specific impacts on *Msm* cell division and cell integrity in real-time, indicating that the phage infection is asynchronous, with the interactions between the phage and host suggested to be impacted by cell state and microenvironment, and that phage infection is associated with increased myco-membrane permeability.

**Figure 2:**
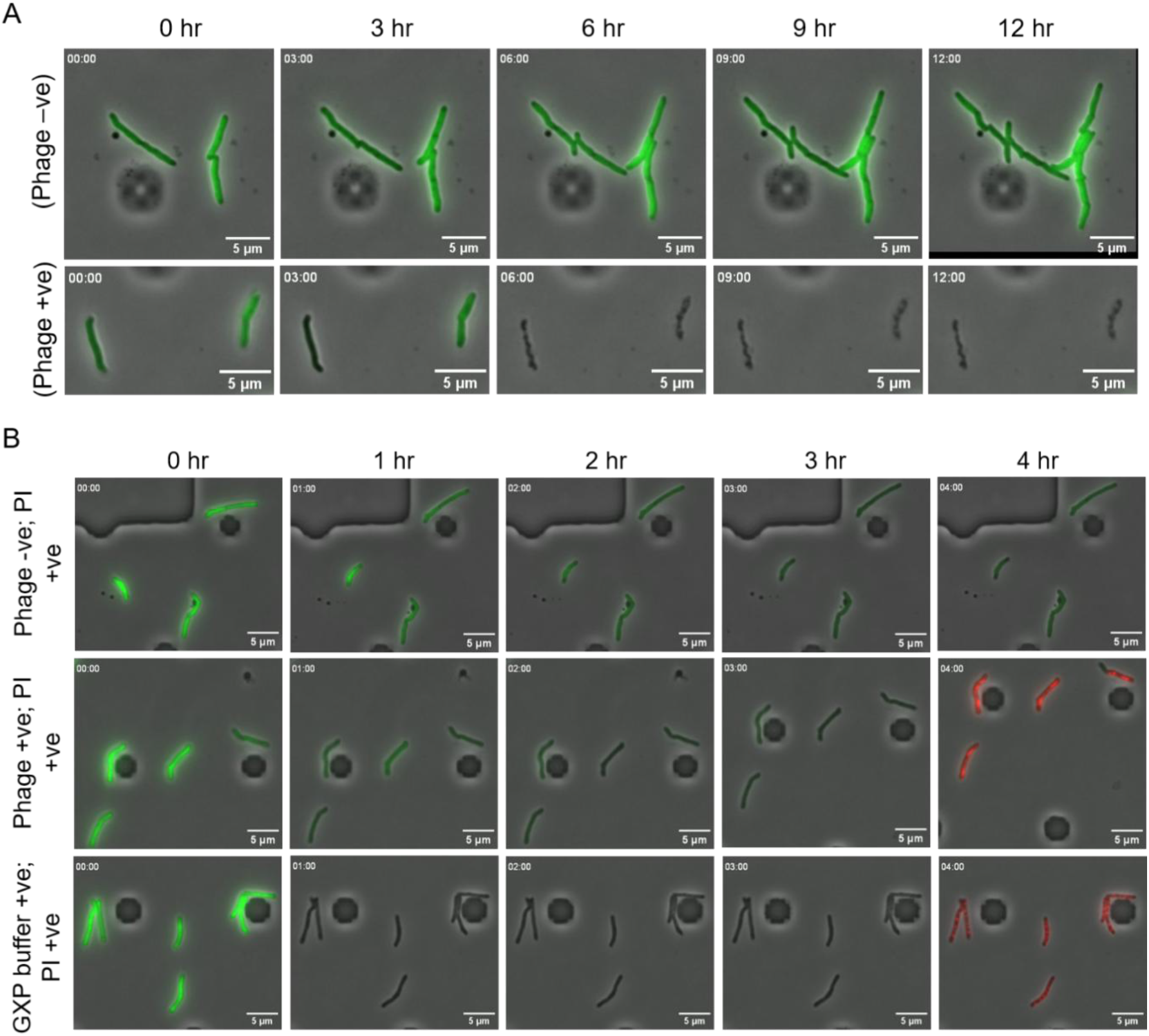
Mycobacterial phage D29 infection drives progressive loss of cytoplasmic contents and mycobacterial permeabilization. (A) A CellASIC perfusion experiment showing representative live-cell time-lapse images captured at 3 hr intervals. The no phage control (top panel) shows mycobacterial division and maintenance of the green fluorescence over the 12 hr window whereas phage infected (phage +ve) show GFP signal loss after 6 hr, suggesting GFP release with mycobacterial cell lysis. (B) An additional perfusion experiment shows representative images recorded at 1 hr intervals. Cells not exposed to phage retained the GFP signal over the 4 hr window but did not take up PI on perfusion (4 hr). In contrast, phage infected cells show loss of cell membrane integrity and mycobacterial viability as evident by uptake of PI at 4 hr post phage infection. As a permeabilization control cells were exposed to GeneXpert (GXP) buffer followed by PI exposure at 4 hr.

### Phage D29 lyses and kills mycobacteria efficiently

In the context of bacterial DNA extraction in this work, we refer to cell lysis as the disruption the bacterial cell envelope sufficiently to release intracellular contents, including gDNA, into the extracellular solution. Notably this lysis may not always require the cell to visibly “burst.” Operationally, a cell is lysed when its envelope has lost its ability to retain the DNA. While productive phage D29 infection results in increased cell permeability and apparent cell lysis with changes in single cell morphologies (transition to ‘ghost’ cells), this is not definitive evidence as DNA may still be contained within cells. To confirm disruption of the mycobacterial cell envelope structures we leveraged propidium iodide (PI), a membrane-impermeant dye that fluoresces red when intercalated into DNA. This selective marker only fluoresces in cells that have membrane disruption and is conventionally used as a marker of cell death (Pooley et al., 2016).

In this experiment, the GFP signal was maintained in mycobacterial controls that were not exposed to phage, and no indication of PI signal was evident in these (**Figure 2B, Supplementary Video 3**). The control (GeneXpert buffer which is used as a proxy to compromise the cell envelope) showed loss of GFP within the first hour of exposure and uptake of PI to indicate cell membrane permeabilization and cell death as reported in a related study (Helb et al., 2010). In phage exposed samples, the loss of GFP signal was observed after 3 hr, and perfusion of PI at 3.5 hr (**Supplementary Methods Figure SM1B**) confirmed mycobacterial membrane disruption indicating cell death (**Figure 2B, Supplementary Video 4**). These results confirmed that the loss of GFP signal observed in the previous experiment (**Figure 2A**) resulted from the loss of cell envelope integrity through phage-mediated permeabilization and membrane disruption resulting in mycobacterial lysis and death.

These single-cell experiments indicate that under these conditions no mycobacterial lysis/ death occurred before 3 hours. In addition, phage lysis was not synchronous with lysis occurring at varying times between 3-8 hr post phage perfusion. This single cell analysis reveals initial lysis events do align with the latent period findings of about 2.5 hr in cell culture assays (**Supplementary Figure S4C**); however, the heterogeneity indicates that population level behaviours in batch culture may not be reflected in the micro-environment of the CellASIC ONIX2 device. This may be due to factors such as the continuous flow of the phage particles over the trapped bacteria during the experiment, which may impede adsorption/ infection, or the dynamics of perfusion which may result in initially low densities of the phage solution in the mycobacterial chambers which delay or cause variability in phage adsorption timing. Based on these findings, a conservative approach involving extended exposure of mycobacteria to phage, in well mixed culture, is recommended for DNA extraction protocols to ensure maximum lysis. During optimization (**Supplementary Figure S4; Supplementary Methods**) and characterization of phage infections, we established a conservative incubation period for effective phage lysis of *Msm*. A 6-hr incubation was selected as this is post the latent period (∼ 150 min), and within the rise period, where a linear increase in phage numbers was observed up to 5 hours (**Supplementary Figure S4C**). This corresponds to approximately two *Msm* generation times (∼3 hr each).

In addition to heterogeneity in infection dynamics, single-cell experiments revealed that not all cells are infected with phage under these conditions. We examined bulk lysis, optimising conditions to characterize efficacies of both phage adsorption and bacterial survival. Fluorescence microscopy of *Msm*::GFP cells stained with SYTOX Orange after phage exposure revealed low numbers of intact cells after phage contact (**Supplementary Figure S5A**). Flow cytometry of *Msm* infected with SYTOX orange labelled phage confirmed that phage adsorption signals varied across the population with 93% of cells SYTOX Orange positive (**Supplementary Figure S5B**). *Msm* WT survival post phage D29 infection, estimated using colony forming units (CFU), was significantly reduced, with <100 cells surviving phage exposure from starting populations of approximately 10^8^ *Msm* bacilli (**Figure 3A**), which represents >99.9999% loss of viability. Examination of a selection of the surviving *Msm* colonies revealed that the majority (8 out of 10 randomly selected and tested) were resistant to phage D29 lysis (**Supplementary Figure S6**). We extended this analysis to *Mtb* H37Ra which demonstrated a comparable loss of viability following phage exposure (**Figure 4A**).

**Figure 3:**
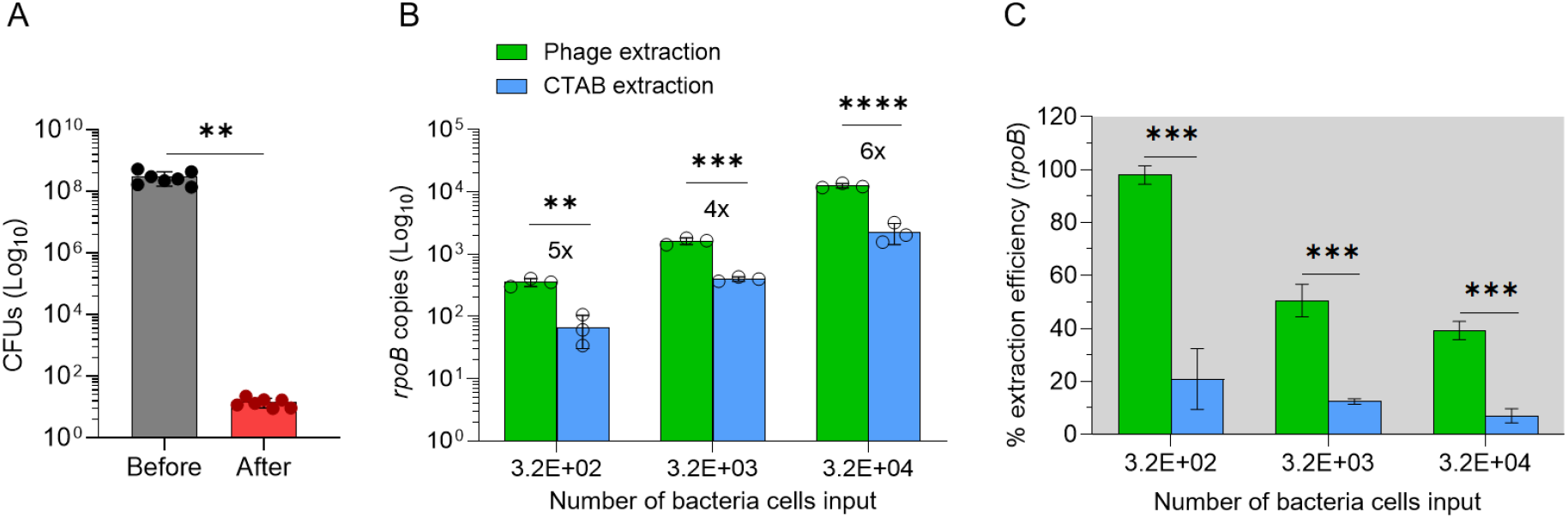
Mycobacteriophage D29 achieves high lysis efficiency and superior gDNA recovery in *Msm* compared to CTAB extraction as quantified using ddPCR. (A) CFU survival of *Msm* that was not exposed to phage (before) *versus* phage treated cells (after). Dots represent independent experiments. Error bars show means with standard deviations, P-values were acquired by paired t-tests. *\*\*P* < 0.01. (B) Quantification of *Msm rpoB* copy numbers in gDNA isolated by either phage D29 (green) or CTAB (blue) methods across three cell densities shown on the x-axis. Fold increase of copy numbers in phage D29-based compared to CTAB extractions are shown above the bars. (C) Efficiencies of the phage and CTAB DNA extractions are calculated relative to the theoretical genome copies in samples containing expected bacterial numbers shown on the x-axis. The data are shown as means. Error bars represent the standard deviations of three biological repeats. The P-values were calculated using unpaired t tests. *\*\*P <* 0.01, *\*\*\*P <* 0.001, *\*\*\*\*P <* 0.0001

**Figure 4:**
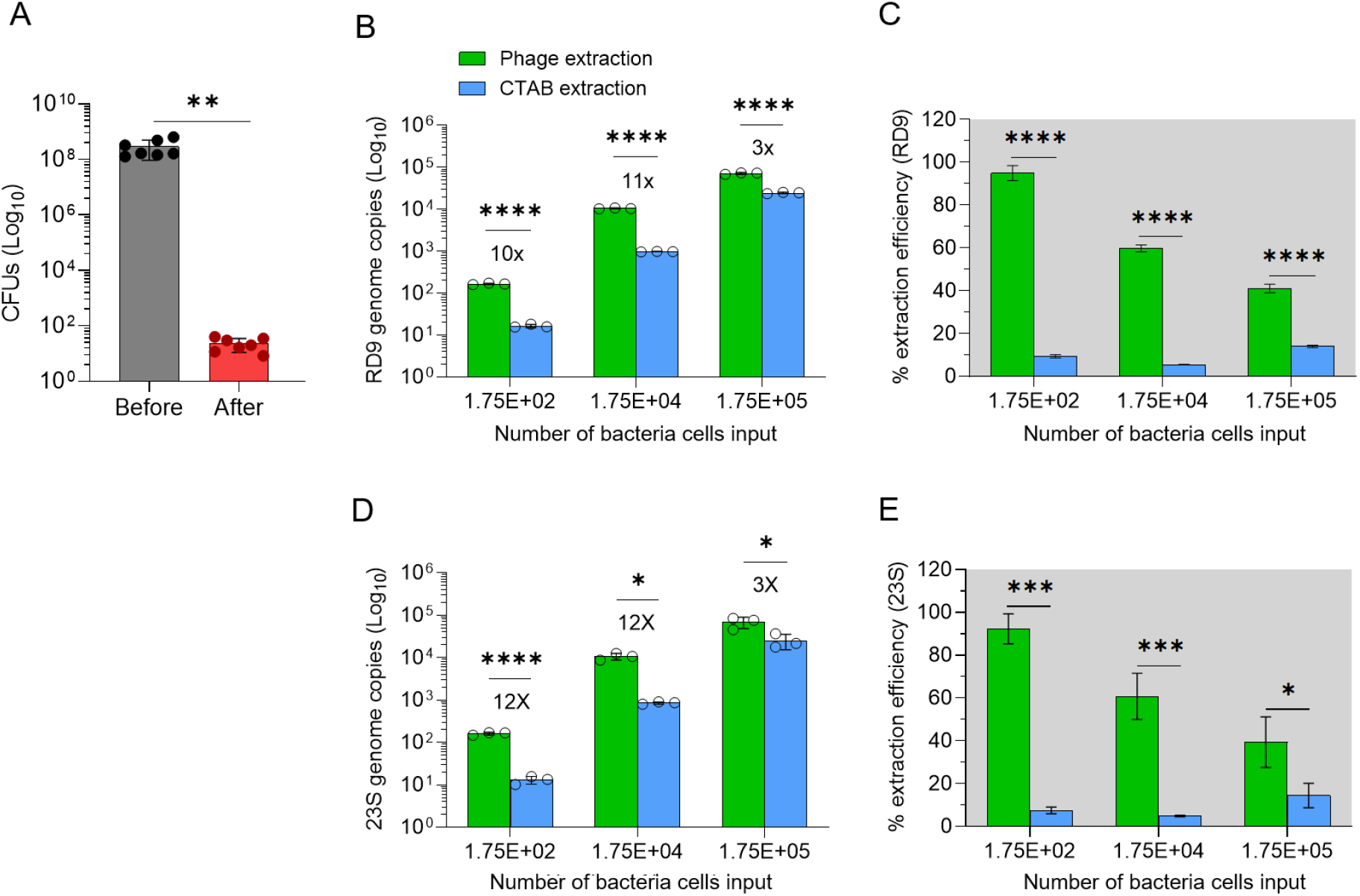
Mycobacteriophage D29-based lysis and DNA extraction efficiency improvements translate to *Mtb*. (A) CFU survival of *Mtb* H37Ra that was not exposed to phage (before) *versus* phage infected samples (after). Dots represent independent experiments. Error bars show means with standard deviations, P-values were acquired by paired t-tests. *\*\*P* < 0.01. (B) RD9 genome copies in gDNA isolated by phage D29 (green) and CTAB (blue) methods across three cell densities shown on the x-axis. Fold increase in yield in phage D29-based compared to CTAB isolations are shown above the bars. (C) Efficiencies of the phage and CTAB DNA extraction calculated relative to the theoretical genome copies of RD9 samples containing bacterial numbers shown on the x-axis as a comparison. (D) genome copies of 23S rRNA and (E) efficiencies of 23S rRNA illustrate reproducibility of approach. The data is shown as means. Error bars represent the standard deviations of three biological repeats. The P-values were calculated using unpaired t tests. *\*P <* 0.05, *\*\*\*P <* 0.001, *\*\*\*\*P <* 0.0001

### Phage D29-mediated lysis enables high-efficiency DNA recovery from low biomass mycobacterial samples

Having established that phage D29 efficiently kills and permeabilises mycobacteria, we next asked whether this lysis translates into efficient recovery of gDNA. The optimized phage lysis conditions (**Supplementary Figure S4**) were applied to quantify *Msm* DNA recovery with CTAB DNA extraction protocol performed in parallel for comparison of yield (**Figure 3B, C**). DNA was quantified first by ultraviolet-visible absorption spectroscopy (Nanodrop) (**Supplementary Figure S7B**) and fluorometry (Qubit fluorometer) (**Supplementary Figure S7C**), and by amplification-based readouts (qPCR and ddPCR). Amplification based quantification of *Msm* WT gDNA targeted *rpoB* (**Figure 3B, C**) and, in *Msm*::GFP, *gfp* (**Supplementary Figure S7D, E**). All targets are single copy in their respective mycobacterial genomes; so, one copy detected was taken as equivalent to one bacterium and this relationship was used to determine extraction efficiency. Targeting *gfp* offered a further advantage, as the *gfp* gene is absent from the WT *Msm* used to amplify phage stocks, quantification of *gfp* therefore reports DNA yield with no possible contribution from the remaining mycobacterial DNA in phage stock DNA.

Nanodrop and Qubit quantification of total gDNA from identical WT *Msm* samples gave 3-to 4-fold higher yields with the phage method compared to CTAB (**Supplementary Figure S7B, C**). By qPCR (*gfp*) and ddPCR (*rpoB*), phage-based extractions yielded consistently higher DNA (4- to 7-fold) than CTAB across the three cell dilutions tested, with statistically significant differences between the two methods (**Figure 3B, C; Supplementary Figure S7D, E**). At low bacillary numbers of *Msm* (100 and 320 cells by qPCR and ddPCR respectively), the phage method achieved approximately 95% extraction efficiency, a marked improvement over the ≤20% efficiency achieved by CTAB using the same inputs (**Figure 3B, C; Supplementary Figure S7D, E**), confirming its suitability for DNA extractions at low bacterial densities. Phage-based extraction efficiency declined as cell density increased, suggesting a decline in phage-based lysis or additional DNA losses during higher biomass processing. CTAB efficiency likewise declined with increasing cell density, indicating a limitation of this method for high density samples. Overall, phage D29-based lysis enabled more efficient gDNA recovery than CTAB in *Msm*, establishing its suitability for low density mycobacterial samples. We note potential application of the phage-based DNA isolation in antibiotic resistant mycobacteria cells through lysis and plaque formation on lawns of rifampicin resistant *Msm* mutant (**Supplementary Figure S4A**).

Although phage-based extraction has utility in *Msm*, its most valuable application will be in pathogenic mycobacterial strains such as *Mtb*. *Msm* and *Mtb* differ in cell envelope complexity, in mycolic acid composition and acid-fastness (Yamada et al., 2018), in the number of L,D-transpeptidases that form peptidoglycan (PG) crosslinks (Baranowski et al., 2018), and in envelope thickness (Daffé & Marrakchi, 2019). These differences could influence susceptibility to phage-mediated envelope disruption and therefore limit the direct translation of the approach from *Msm* to *Mtb*. We therefore tested the phage lysis method in the attenuated strain *Mtb* H37Ra. We infected approximately 10^8^ CFUs/mL with phage at multiplicity of infection (MOI) of 100 and first assessed CFU (**Figure 4A**) and then quantified phage-based and CTAB DNA recovery by ddPCR targeting the region of difference 9 (RD9) and 23S rRNA gene. Consistent with *Msm*, phage D29-based isolations yielded higher gDNA copy numbers than CTAB, by both RD9 (**Figure 4B, C**) and 23S rRNA (**Figure 4D, E**) across all cell densities tested. At the lowest input (175 cells/mL) phage-based efficiencies exceeded 92%, with CTAB 10- to 12-fold lower (below 10% extraction efficiency). Differences in DNA yield were statistically significant at every input. As observed in *Msm*, phage D29-based efficiency declined at higher bacterial numbers. CTAB efficiency, by contrast showed a slight decline and then an improvement as bacterial densities increased from 175 to 17500 to 175000, but remaining below 20% extraction efficiency across this cell density range.

Together, these findings show that the phage D29-based DNA extraction is an improved alternative to CTAB for DNA isolation in both *Msm* and *Mtb*. Notably, the phage-based extraction efficiency increased with decreasing bacillary density, underscoring its potential for sensitive molecular applications on low biomass samples.

## Discussion

Efficient and effective lysis of bacterial cells is a prerequisite for recovering intracellular molecules, including gDNA, for downstream molecular applications ranging from diagnostic detection to whole genome sequencing and population level genomics. In mycobacteria, this is a particular challenge: the exceptionally complex cell envelope, characterized by an outer mycolate layer linked to an arabinogalactan-peptidoglycan scaffold (Jacobo-Delgado et al., 2023), confers both mechanical rigidity and chemical impermeability that render conventional lysis approaches inefficient (de Bruin et al., 2019). Here we show that phage D29-based lysis substantially improves gDNA recovery over the standard CTAB method and, using single-cell live imaging, we resolve features of this bio-technical application. This work expands on our understanding of the parameters governing phage infection efficiency including phage exposure duration, MOI, adsorption sites and kinetics.

Phage adsorption is the entry point for infection with the presence and localisation of phage reporting on the features and states of the mycobacterial cell envelope that permit attachment. Phage D29 adsorbs preferentially at the poles and septa of *Msm*, visualized through SYTOX Orange labelling of the phage. The polar signal was frequently unbalanced between opposite poles, or present at only one pole, indicating a preference for adsorption at one pole. Using the 4-N,N-Dimethylamino-1,8-naphthalimide conjugated trehalose (DMN-Tre) probe (Kamariza et al., 2018), we confirmed the differences in mycobacterial polarity in our strains and conditions which arise from differential elongation between the older ‘mother’ poles and newer ‘daughter’ poles, consistent with other studies (Dinkele et al., 2021; Maitra et al., 2019; Rimal et al., 2022; Thanky et al., 2007), and with observations using a N-quencher trehalose fluorophore (N-QTF) probe (Dulberger et al., 2023). Together, the SYTOX Orange profiles indicate that phage D29 adsorbs at the poles and septa, the zones of active elongation and *de novo* PG biosynthesis (Maitra et al., 2019; Rimal et al., 2022; Thanky et al., 2007). Dulberger et al. (Dulberger et al., 2023), reported analogous polar/septal attachment for bacteriophages (namely Fionnbharth, Muddy, and Adephagia) on N-QTF labelled *Msm*, whereas polar and septal adsorption pattern was not clearly demonstrated for the mycobacteriophage Bxb1 (Freeman et al., 2025), so the universality of this pattern in mycobacteriophage attachment is not established. Although our study did associate phage attachment with the new and old poles, our data provide additional evidence of growth asymmetry under the environmental conditions tested that is aligned with the preference for phage D29 adsorption at the zones of nascent PG biosynthesis.

Live-cell imaging revealed heterogeneity in timing of phage-induced permeabilization (as demonstrated by loss of GFP and disruption of cell morphology. Individual cells lysed asynchronously, and not all cells were infected under the conditions tested, revealing variation that population-level assays can’t resolve. This variation may be, in part, technical, possibly reflecting non-uniform phage-cell contact under continuous perfusion in the CellASIC device leading to lack of adequate attachment; however, this does expose heterogeneity and reveal that multiple factors including bacterial state, microenvironmental conditions as well as phage-cell ratios will impact infection rates and outcomes. These infection dynamics informed the phage-exposure times used for DNA extraction. As the mycobacteriophage lytic cycle scales with host generation time (McNerney et al., 2004), and the earliest single-cell lysis events (∼3 hr) approximated the *Msm* latent period established in this study (∼150 min), and generation time (∼3 hr;) (Ning et al., 2021), we adopted a conservative 6 hr phage-exposure window for *Msm*. This is comparable to the timing determined for the Actiphage system (Swift et al., 2020), where a 120 min latency period was estimated. For the slow-growing *Mtb* H37Ra (doubling time of ∼18-24 hr), we scaled the window to 24 hr. We note that this *Mtb* window was inferred from an approximate generation-time scaling rather than optimised empirically and represent an assumption to be refined in future work. Both windows extend the 3 hr incubation time established in the Actiphage method (Swift et al., 2020).

Across all quantification approaches, phage D29-based extraction was superior to CTAB in both *Msm* and *Mtb*. Strikingly, at very low cell numbers the phage-based method recovered close to the theoretical DNA yield, indicating near-complete lysis, with minimal downstream loss. This quantitative performance extends the phage-diagnostic lineage: phage D29 has been used as a lysis agent to detect mycobacteria in cattle from samples containing as few as <10 (Swift et al., 2020) and <100 bacteria (Swift et al., 2016). It has been also been used in the diagnosis of *Mtb* infections from human blood samples (Verma et al., 2020). While these studies establish detection in the veterinary or clinical context, our data quantify recovery efficiency and benchmark it directly against a standard method across a range of cell densities. CTAB performed poorly at low density bacterial samples, where sample losses and/ or incomplete lysis likely dominate, implying it is unsuitable for paucibacillary material. Because expanding such samples in cultures risks altering the sample composition, direct phage-based DNA extraction offers a route to interrogate low-biomass populations without the culture-adaptation bias associated with in vitro propagation, though the method introduces its own selectivity toward phage-susceptible cells whose implications for rare-variant recovery remain to be systematically evaluated.. The practical application of the phage-based extractions to paucibacillary clinical samples such as blood or bioaerosol *Mtb* samples (Barr et al., 2021; Swift et al., 2020), which potentially contain unique and rare genotypes of interest, is recommended. Our single-cell characterization of phage attachment and lysis illustrate that this approach is suited to be paired with microfluidics technologies, to enable low biomass sample handling, with future downstream molecular applications such as nanopore sequencing.

The main limitation observed was a decline in recovery efficiency at high cell density: although absolute yields remained higher than CTAB, they fell below theoretical expectations as bacterial numbers increased. We interpret this primarily as a recovery step ceiling rather than a failure of phage lysis. As near-complete lysis (>6-log₁₀ loss of viability by CFU) was achieved at ∼10⁸ cells (a density higher than any used in the DNA extraction efficiency assays), incomplete phage attachment or lysis is unlikely to account for the fall in recovery efficiency at high cell number, implicating loss at the DNA recovery step (possibly retention on the debris filter or saturation of the clean-up column) rather than a failure of phage killing. A biological contribution cannot be excluded; other mechanisms such as abortive infection may play a role in “death without lysis” owing to activation of processes such as programmed cell death (Calcuttawala et al., 2022; Samaddar et al., 2016). This would leave cell envelopes intact and DNA unreleased. Importantly, the phage-based method still outperformed CTAB at every density and excelled in the low biomass regime. The high-density gap remains to be explored, with the first step being to examine this as an efficiency metric artifact of near saturation of downstream extraction steps rather than a limitation of phage lysis itself.

A limitation of this study is that quantitative superiority was only benchmarked against CTAB extraction. Mechanical methods (bead-beating), commercial kits, and hybrid protocols such as the recently described chloroform-bead method (Murase et al., 2025) were not evaluated, and the fold-improvement magnitudes reported here should be interpreted specifically in the context of comparison against CTAB.

The phage-based method is inherently selective for viable, phage-susceptible mycobacteria. This feature that enables selective detection of live cells but also implies that phage-refractory subpopulations may be underrepresented in recovered DNA. We observed rare *Msm* survivors of phage exposure whose lawns were not impacted by phage spotting assays, indicating phage resistance. Resistance may be transient or genetically stable, for example, overexpression of the multicopy phage-resistance (*mpr*) gene alters the *Msm* envelope and blocks productive injection by phage D29 and its close relative, phage L5 (Barsom & Hatfull, 1996). Although rare in our hands, such resistance sets a ceiling of the completeness of lysis and warrants further study, in particular what the mechanisms conferring resistance in these *Msm* colonies are and whether analogous mechanisms operate in *Mtb*. Whether this selectivity introduces a systematic bias against rare genotypes of clinical interest (for example, drug-resistant variants) depends on whether phage susceptibility co-segregates with these genotypes. This question is not addressed by the present study and represents an important prerequisite for applying phage-based extraction to microheteroresistance detection and mixed-strain sampling.

## Conclusions and outlook

Phage-based lysis is a gentle, species- and viability-selective means of compromising the mycobacterial cell envelope with a biological agent. Its extraction efficiencies for low input samples are suited to paucibacillary clinical samples and its activity against a rifampicin-resistant *Msm* mutant suggests applicability to drug-resistant strains, for which rapid, culture-independent genomic access is urgently needed (Singh et al., 2023). The single-cell phenotypes uncovered here, including heterogeneous adsorption and asynchronous lysis, argue for interrogating mycobacterial populations at low density and at the level of the individual cell, where single-cell measurements can expose the variation that underlies population-level behaviours and outcomes (Chung et al., 2024). The ability to follow phage infection thorough perfusion in a microfluidic device point to a natural extension of this work in marrying microfluidic single-cell handling with phage-based molecular diagnostics. Pairing this with short and long-read sequencing (Hall et al., 2023) to enable recovery of clean, native DNA with subsequent sequencing of individual bacterial cells, would enable direct study of paucibacillary clinical material such as blood, bioaerosol and tongue-swab samples (Barr et al., 2021; Dinkele et al., 2021; Swift et al., 2020). Beyond DNA extraction, these findings point to a broader agenda in precision infectious-disease diagnostics (Woodhouse et al., 2024) with the same envelope-compromising activity of phage D29 harnessed for more efficient and selective cell labelling, enhanced antibiotic access and viability assessment.

## Experimental Procedures

### Mycobacterial strains and culture conditions

Key resources and mycobacterial strains used in this study are listed in **Supplementary Methods, Table S1, Table S2 & Table S3**. Unless otherwise stated, mycobacteria strains were cultured in Middlebrook 7H9 supplemented with 0.2% glycerol, 10% ADC and 1 mM CaCl_2_, without Tween 80 (phage medium). Liquid cultures were incubated at 37°C with shaking at 150 rpm. Solid cultures were grown on Middlebrook 7H10 supplemented with 0.5% glycerol, 10% ADC and CaCl_2_ to a final concentration of 1 mM and maintained at 37°C. Stocks were stored at -80°C in 33% glycerol.

### Mycobacterial enumeration

Bacterial densities were determined by FC using SYBR Gold staining, as previously described (Barr et al., 2021). Briefly, cultures (OD_600_ = 0.3-0.4) were pelleted and resuspended in an equal volume of phosphate buffered saline solution plus 0.1% Tween 80 (PBST). These were ten-fold serially diluted in PBST, needle-dispersed, and heat-permeabilized. Sampling was performed on BD Accuri C6 flow cytometer and analysis performed using FlowJo. Mycobacterial viable counts as CFUs were determined, where indicated by plating serial dilutions on Middlebrook 7H0 with standard supplementation as previously described (Chengalroyen et al., 2024).

### Phage amplification and titration

Phage D29 was amplified in WT *Msm* cells (mc^2^155) using the soft agar overlay method (Hyman & Abedon, 2009; McNerney et al., 2004; Rees & Botsaris, 2012). Briefly, 200 µL of exponential phase (OD_600_ = 0.3-0.4) bacteria, in phage medium, were infected with diluted phage stocks containing approximately 10^3^ phage particles and incubated at 37°C for 30 min to allow phage adsorption. The phage-adsorbed bacteria were incorporated into soft agar and overlaid onto 7H10 agar. After 12 hr of incubation at 37°C the phage particles were recovered in phage buffer filtered through 0.22 µm membranes and treated with DNase I to remove contaminating bacterial DNA and then kept at 4°C. Phage titers were determined by serial dilution and titers expressed as PFU/mL (Bichet et al., 2021).

### Multiplicity of infection and bacterial survival

Exponential phase (OD_600_ = 0.3-04) *Msm* were either left uninfected (control) or infected with phage particles at estimated MOI of 1, 10, 100 and 1000. Control samples were plated for CFU immediately while infected samples were plated for CFU following incubation for 12 hr at 37°C. The plates were incubated at 37°C for 5 days and CFUs counted with bacterial survival of infected cells expressed relative to the uninfected controls.

### Phage latent-period determination

Phage D29 latent period was determined using a modified phage-amplification assay (Botsaris et al., 2010; McNerney et al., 2004). Briefly, exponential phase *Msm* cultures (OD_600_ = 0.3 which is estimated as ∼10^5^ CFU/mL) were infected with approximately 10^5^ phage particles (MOI = 1) and incubated at 37°C for 30 min. Following phage adsorption extracellular phage was inactivated with 10% of 100 mM ferrous ammonium sulphate (virucide), allowing a contact time of 5 min. Cells were washed and resuspended in fresh medium and incubated at 37°C with shaking at 150 rpm. This culture was sampled at 30 min intervals for phage titration by plaque assay as described previously (Bichet et al., 2021).

### Fluorescent labelling and imaging of phage adsorption

High-titer phage stocks were stained using a previously described protocol (Van Valen et al., 2012). Briefly, phage D29 particles were labelled with SYTOX Orange (500 nM) and incubated for 3 hr at room temperature. Free stain was removed by washing the samples four times in phage buffer using Amicon Ultra-0.5 mL Centrifugal filters Ultracel®-30K. Stained phage particles were recovered using phage buffer, titrated and used within one week of staining. The fluorescent reporter strain, *Msm*::GFP (Chan et al., 2002) (OD_600_ = 0.3) was infected with the labelled phage at an approximate MOI = 0.01 and incubated for 30 min. Cells were washed and imaged using a Zeiss Axio Observer 7 microscope equipped with a 100X 1.4 NA plan apochromatic phase 3 oil immersion objective lens. Image capture was performed on the Zeiss ZEN software (Zeiss). GFP was excited using a GFP detector (excitation wavelength 450-490 nm and emission wavelength 500-550 nm) and filtered at 450-488 nm, exposure time of 150 milliseconds (ms) was used in the experiment. Red fluorescence (SYTOX Orange) was excited using Alexa Fluorescence 568 detector (excitation wavelength 550-580 nm and emission wavelength 590-650 nm) and filtered at 540-570 nm, exposure time of 150 ms was used in the experiment. Images were captured using Axiocam 506 mono camera (Zeiss). Image processing was performed using Fiji (Schindelin et al., 2012), and fluorescence profiles were analysed using MicrobeJ plugin (Ducret et al., 2016).

### Live-cell time lapse microscopy

Live-cell imaging was performed using a CellASIC ONIX2 microfluidic system with B04A bacteriology plates mounted on the Zeiss Axio Observer 7 system for imaging as described above. Exponential phase mycobacteria cells (OD_600_ = 0.3-0.4) were filtered through 5 µm membranes, loaded into the microfluidic chambers and experimental specific protocols (**Supplementary Methods; Figure SM1**) run to expose cells to the indicated treatments under continuous perfusion. In brief, GFP release, GFP release and uptake of PI and BIODIPY-FL-van assays, were monitored using GFP expressing or mScarlet expressing *Msm* (**Table S2**). For each experiment, images were recorded at defined intervals and analysed using Fiji and MicrobeJ. Detailed experimental protocols (including perfusion protocol and acquisition settings) are provided in (**Supplementary Methods**).

### CTAB and phage D29-mediated DNA extraction

Mycobacterial cells (*Mtb* H37Ra, *Msm* mc^2^155 and *Msm*::GFP) were precultured from stock and sub-cultured twice in phage medium to reach an OD_600_ = 0.4. Cultures were enumerated by FC (Barr et al., 2021) and CFUs. Equivalent bacterial cell inputs were processed using either CTAB method (van Helden et al., 2001) or phage D29-mediated lysis where 1 mL aliquots of bacterial culture were infected with the phage at an estimated MOI = 100 and incubated at room temperature for 30 min. Following adsorption, *Msm* and *Mtb* H37Ra cells were incubated for 6 hr and 24 hr, respectively. Samples were then filtered through 0.22 μm membranes to remove intact cells and debris. The filtrate containing mycobacterial DNA was then purified and concentrated using Zymo DNA Clean and Concentrator kit.

### Quantification of *Msm*::GFP DNA by qPCR

DNA recovered from defined numbers of GFP expressing *Msm* cells, was quantified by qPCR using primers targeting the chromosomally integrated *gfp* gene (**Supplementary Methods Table S3**). Reactions were prepared using the Power SYBR® Green PCR Master Mix and quantification performed against a serially diluted gDNA standard. Cycling conditions and reaction compositions are provided (**Supplementary Methods**). Data were analysed in Excel and GraphPad Prism.

### Quantification of mycobacterial DNA by ddPCR

DNA recovered by CTAB and phage D29-mediated extraction was quantified by ddPCR. *Msm* DNA was detected by targeting the *rpoB* gene and *Mtb* H37Ra DNA was quantified using assays targeting both the 23S ribosomal RNA (rRNA) gene (Walter et al., 2021) and RD9 (Brosch et al., 2002; Patterson et al., 2018). Primers and probes can be found in **Supplementary Methods Table S3** with additional details in **Supplementary Methods.** Reactions were set up using ddPCR SuperMix and Prime time Assay, partitioned on a QX200 Droplet Generator, amplified in a Bio-Rad T1000 thermal cycler and read using QX200 Droplet Reader. Data were analysed using QuantaSoft Software, Excel and GraphPad Prism.

## Data availability

### Underlying raw data

Figshare: https://doi.org/10.6084/m9.figshare.33102155

### Supplementary videos

Figshare: https://doi.org/10.6084/m9.figshare.33102248

## Author contributions (CRediT)

*Joseph Wambugu Gitari*: Conceptualization, Data curation, Formal analysis, Investigation, Methodology, Project administration, Validation, Visualization, Writing – original draft, Writing – review & editing

*Anastasia Koch*: Conceptualization, Funding acquisition, Methodology, Resources, Supervision, Writing – review & editing

*Elizabeth Kigondu*: Conceptualization, Methodology, Supervision, Writing – review & editing

*Digby F. Warner*: Conceptualization, Funding acquisition, Methodology, Resources, Supervision, Writing – review & editing

*Mandy K. Mason*: Conceptualization, Funding acquisition, Methodology, Resources, Supervision, Project administration, Writing – review & editing

## Supporting information

https://doi.org/10.6084/m9.figshare.33102041

## Acknowledgements

We thank Professor Graham F. Hatfull and Deborah Jacobs-Sera (Hatfull Lab, University of Pittsburgh, Pittsburgh, PA, USA) for the gift of mycobacteriophage D29. We thank Dr Melissa Chengalroyen (MMRU, University of Cape Town) for providing the *Msm*::GFP and *Msm*::mScarlet reporter strains. We thank Dharanidharan Ramamurthy (MMRU, University of Cape Town) for the provision of the 23S and RD9 ddPCR probes and primer sets.

