## Supplementary material for "Mycobacteriophage D29-mediated lysis improves recovery of mycobacterial genomic DNA from low-biomass samples": https://doi.org/10.6084/m9.figshare.33102041

### Supplementary Figures

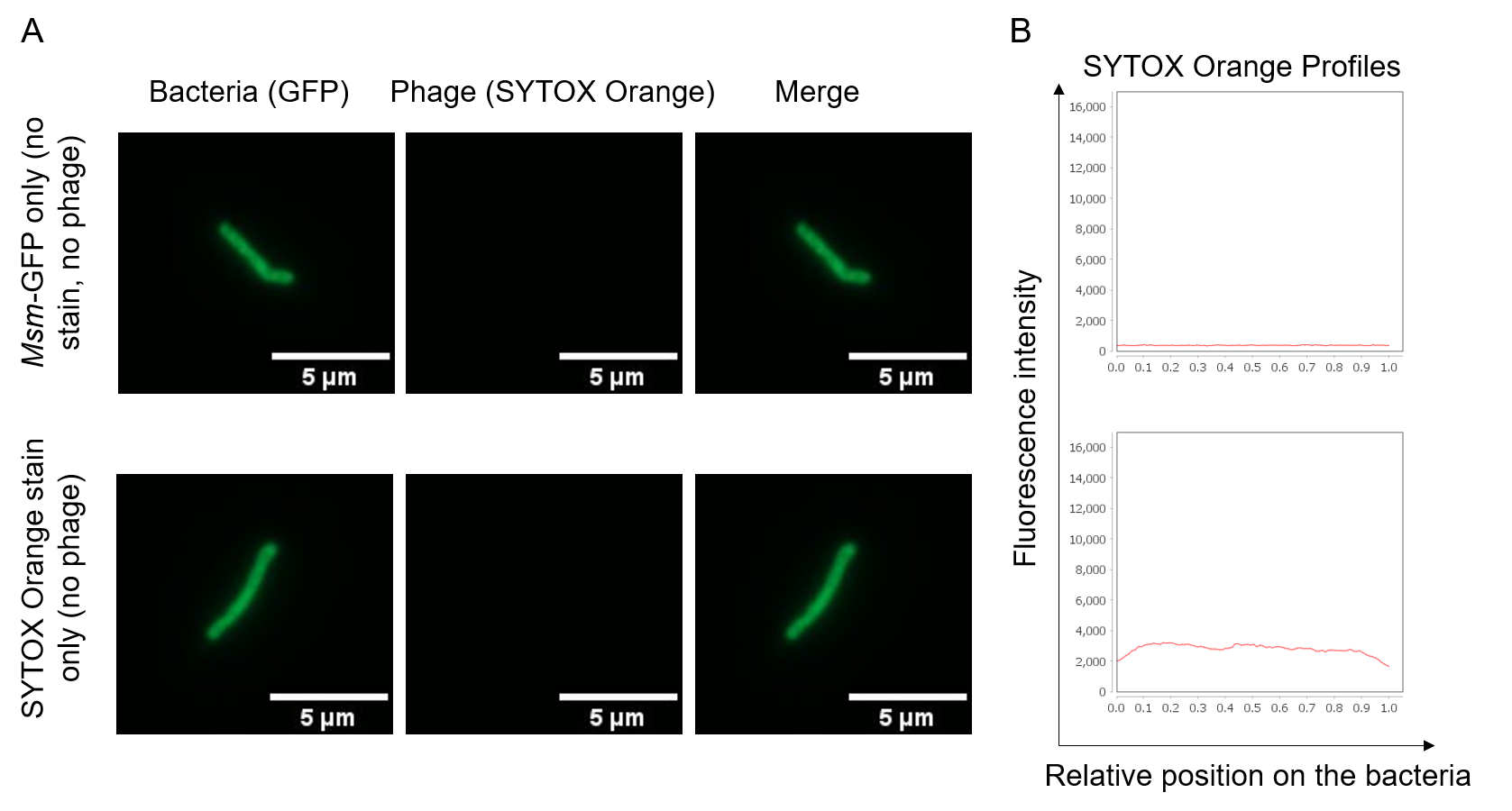

**Figure S1. Control experimental results showing absence or background SYTOX Orange fluorescence signal.** (A) No SYTOX Orange signal was detected when *Msm*::GFP bacteria were imaged in the absence of SYTOX Orange stain and phage D29 or when SYTOX Orange was added to *Msm*::GFP bacteria in the absence of phage D29. (B) Background SYTOX Orange fluorescence signal profiles were observed in the absence of SYTOX Orange stain and phage D29 or when SYTOX Orange was added to *Msm*::GFP bacteria in the absence of phage D29.

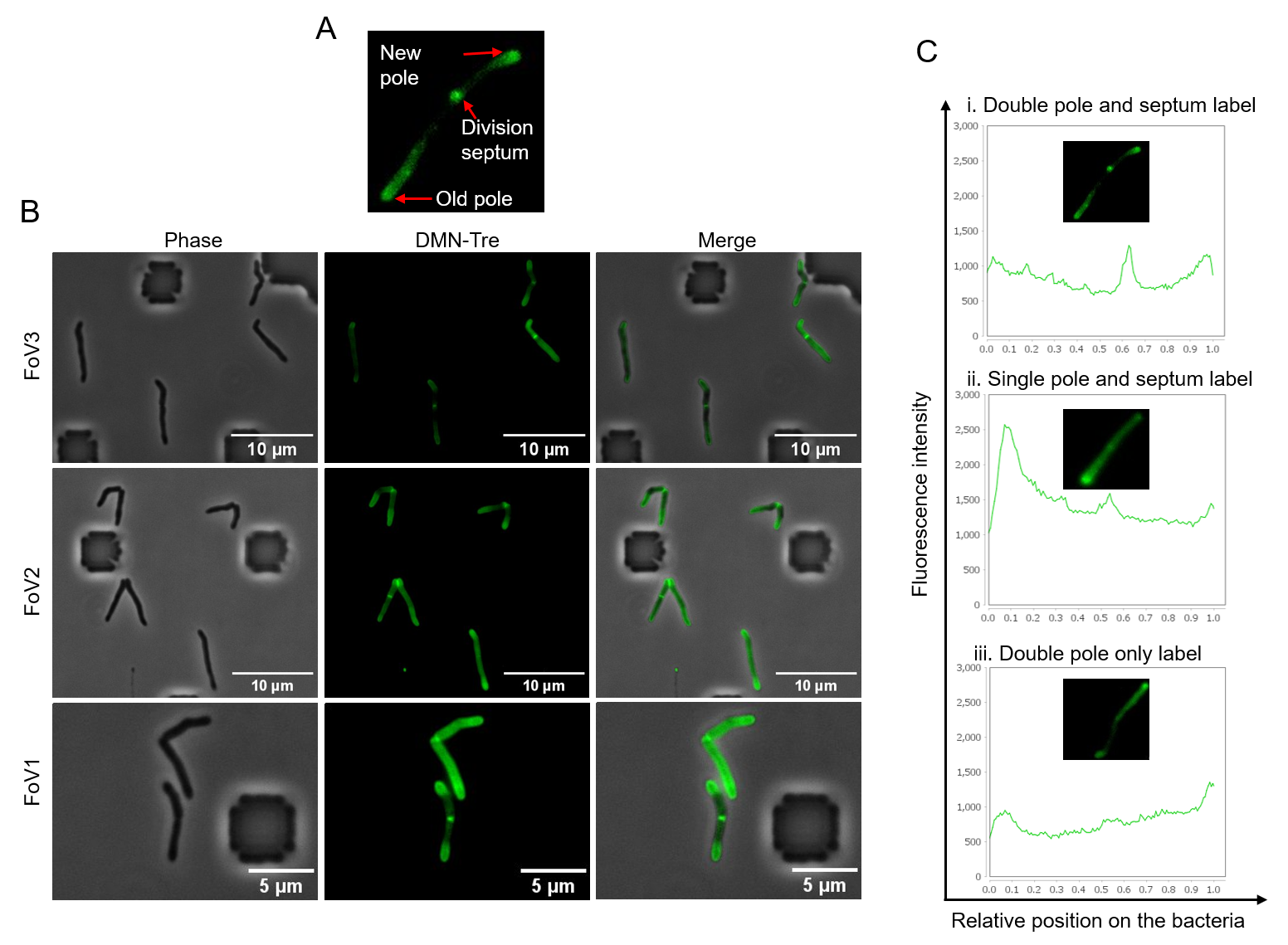

**Figure S2: DMN-Tre labelling of wildtype (WT) *Msm* at the poles and septa.** (A) Incorporation of DMN-Tre at the new pole, old pole, and septum of WT *Msm*. (B) Distinctive DMN-Tre labelling of the poles and septa in separate experiments performed on the CellASIC ONIX2 device. (C) DMN-Tre profiles to the bacterial cell lengths; fluorescence peaks correspond to polar and septal labelling. FoV, field of view

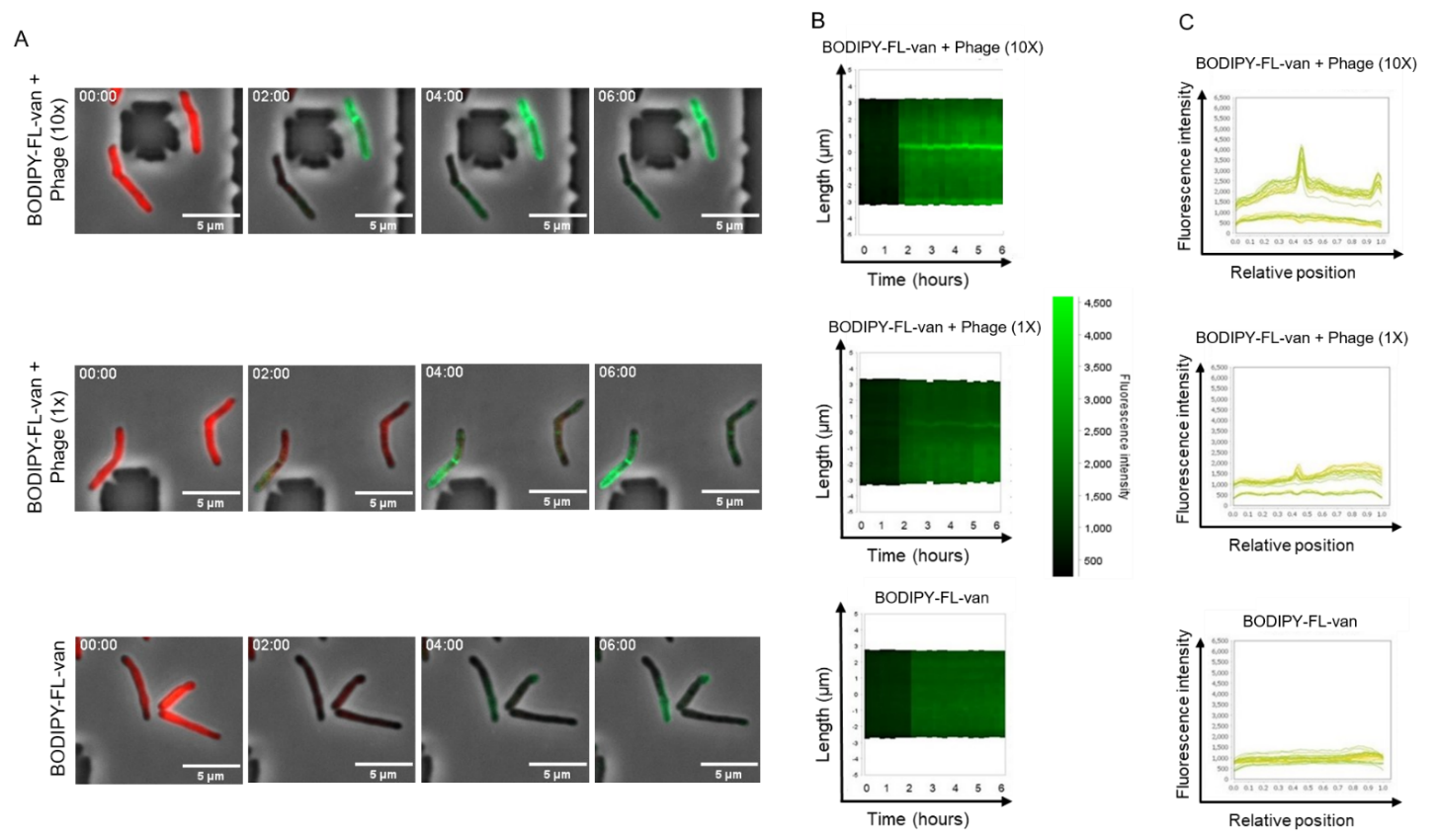

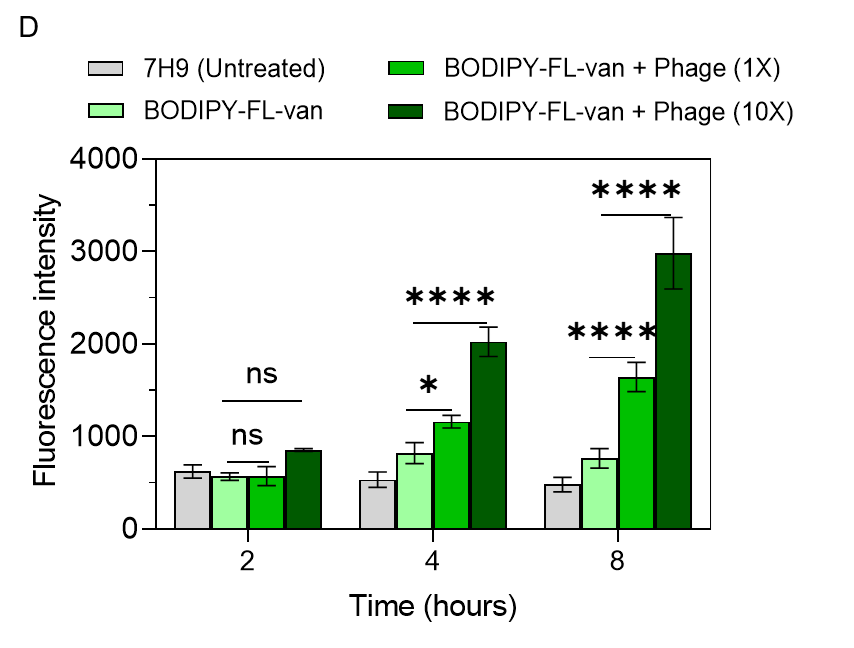

**Figure S3: Mycobacteriophage D29 exposure permeabilizes the cell envelope in a dose-dependent manner.** (A) Live-cell time-lapse images of *Msm*::mScarlet bacteria under high dose – 2.2X10^4^ PFU/mL (10X) of phage D29, low dose – 2.2X10^3^ PFU/mL (1X) of phage D29, and no phage D29 perfusion. BODIPY-FL-van was perfused for 5 hr post phage infection and uptake (green) monitored over time. (B) Kymograph depiction of BODIPY-FL-van labelling (bright green signal) of representative *Msm*::mScarlet cells imaged over 6 hr in the experimental conditions. (C) Profiles of BODIPY-FL-van fluorescence relative to the bacterial cell length show peaks representing accumulation at the poles and septa. (D) Average fluorescence intensity showing accumulation of BODIPY-FL-van over time in a phage concentration-dependent manner. Data shown are the means and standard deviations of 9 representative bacteria. The P-values were calculated using Tukey’s multiple comparisons test after a two-way ANOVA. (*ns P* > 0.05; **P* < 0.05 and *****P* < 0.0001; ns, not significant).

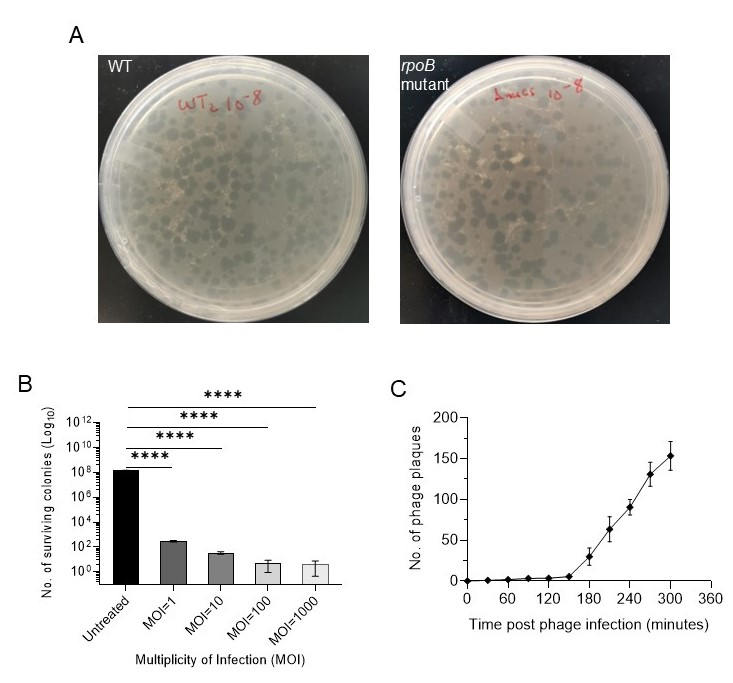

Figure S4: Optimization of phage D29 infection of *Msm*.

(A) Formation of clear phage plaques on lawns of WT *Msm* mc^2^155 and *Msm rpoB* mutant – a mc^2^155 derivative carrying a *rpoB* mutation (codon 442 A->G) conferring rifampicin resistance. (B) Viable bacteria (determined by CFU) post phage infection with increasing multiplicity of infection (MOI). Error bars represent standard deviation derived from four technical repeats. P-values were acquired by Tukey’s multiple comparison tests after one-way ANOVA, *****P <* 0.0001. (C) The number of phage D29-induced plaques determined at 30-minute intervals. The error bars represent the standard deviations of two independent experiments.

The significant reduction in CFU survival from ~10^8^ to ~10^2^ when the MOI was >10 is an indication of highly efficient phage infection, injury and lysis. There was increased cell death with increased MOI, but only a slight reduction in the surviving CFUs at MOI = 1000 compared to MOI=100; therefore, all phage-based DNA extractions (downstream) were performed at MOI = 100 to provide a balance between lysis efficiency while avoiding possible phage saturation and minimize the development of phage resistance.

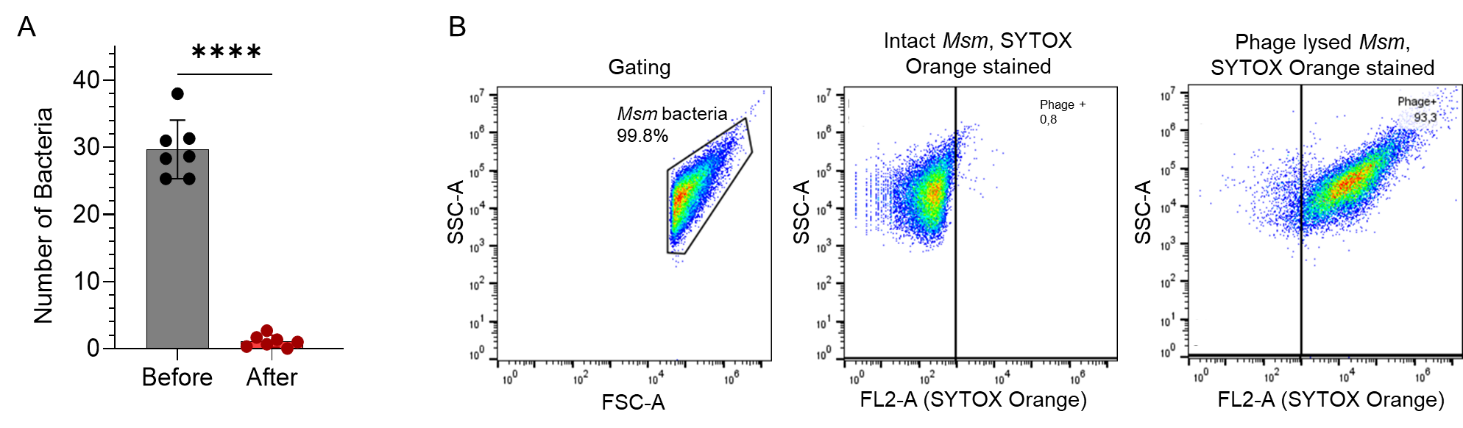

**Figure S5: Quantification of phage D29 lysis of *Msm* cells**

(A) Fluorescence microscopy data showing the number of *Msm* bacteria before and after phage exposure quantified by the uptake of SYTOX Orange, dots represent independent experiments. Error bars show means with standard deviations, P-values were acquired by paired t-tests. *****P* < 0.0001 (B) Flow cytometry plots showing scattering of intact and phage-exposed *Msm* after addition of SYTOX Orange*.* Phage permeabilization and lysis of mycobacteria resulted in the shift of the population to the right quadrant. Flow cytometry was performed on a BD Accuri C6 (488 nm excitation, and emission collected in a 585/40 bandpass filter in the FL2 channel). *Msm* bacteria were first gated on SSC-A/FSC-A before performing the analysis.

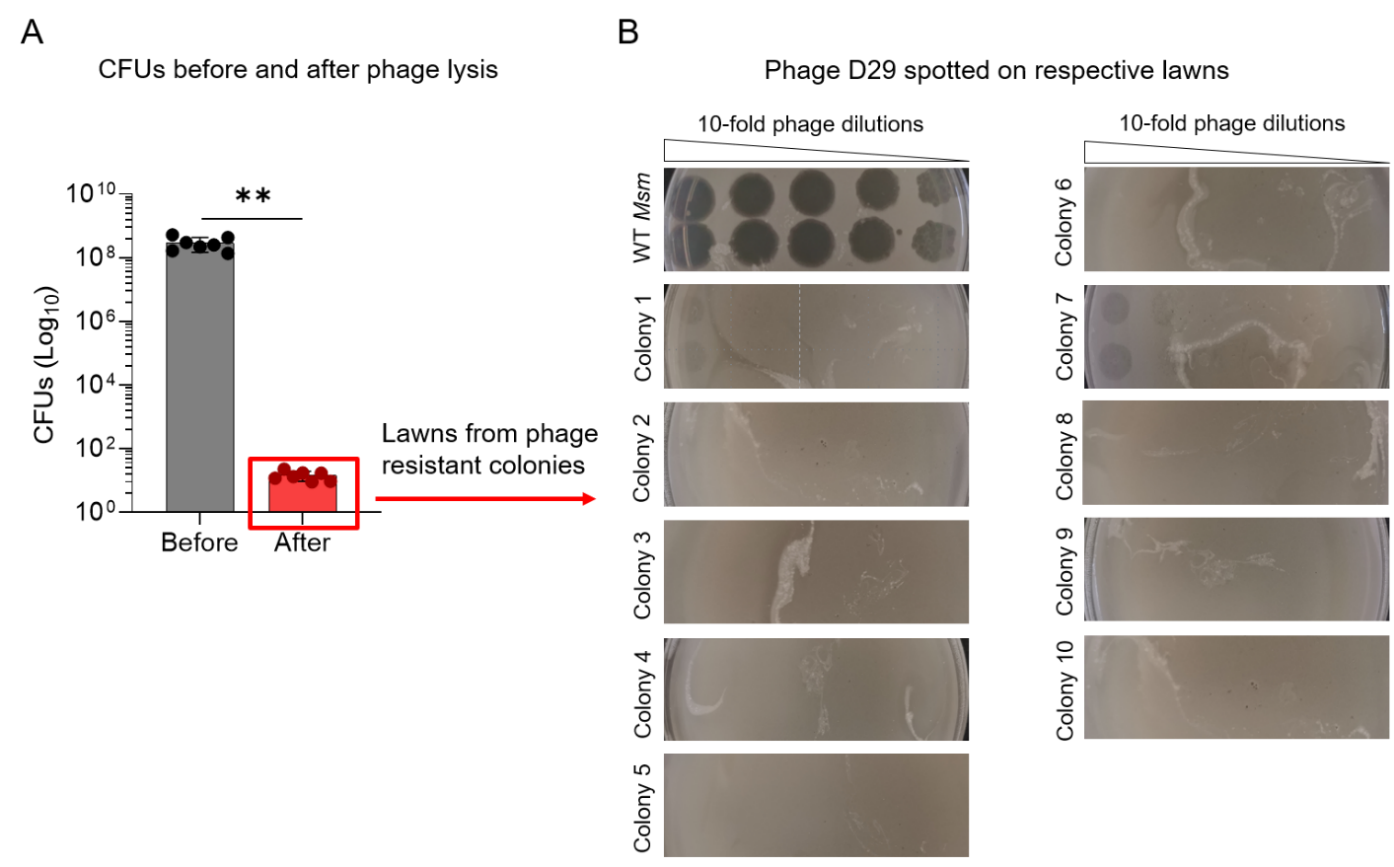

**Figure S6: *Msm* colonies surviving phage exposure show reduced phage susceptibility.** (A) *Msm* survival before (grey bar) and after (red bar) phage D29 infection, with <100 cells surviving from starting populations of ~10^8^ bacilli (adapted from **Figure 3A**). Dots represent independent experiments and bars the means, with error bars showing standard deviations. P-values were acquired by paired t-tests. ***P* < 0.01. Several surviving colonies (boxed in red) were picked and used to prepare lawns for phenotypic assessment of phage susceptibility. (B) Serial dilutions of phage stocks spotted in duplicate on lawns from the phage surviving colonies: WT *Msm* was sensitive, forming clear plaques at all dilutions; two picked colonies (1 and 7) showed slight sensitivity with plaque formation at the highest phage concentration, whereas eight picked colonies (2, 3, 4, 5, 6, 8, 9 and 10) showed a loss of phage sensitivity, with no plaque formation at all phage dilutions. Phage spotting assay data are from a single experiment.

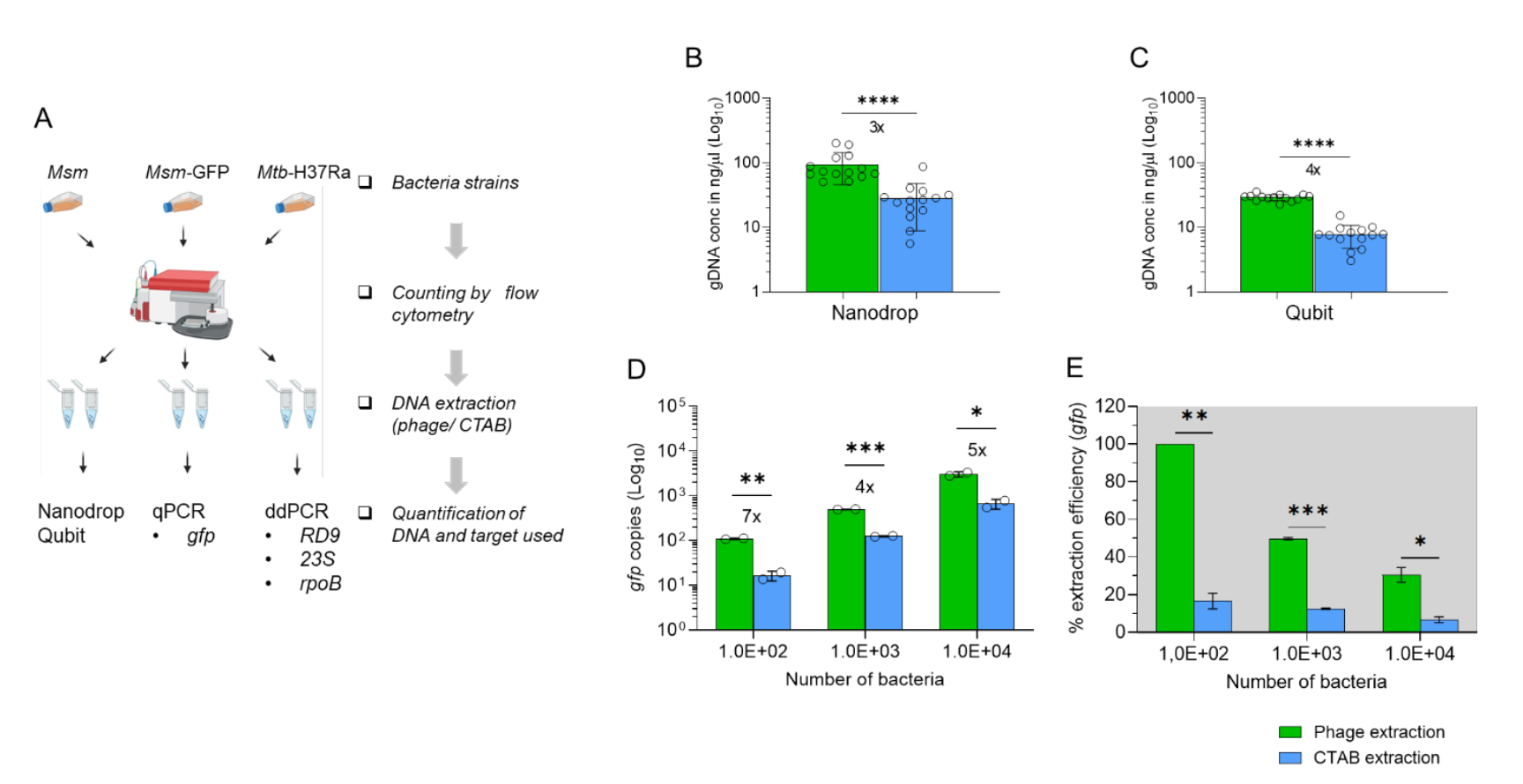

**Figure S7: The isolation and quantification of gDNA in *Msm* - phage D29 lysis *versus* CTAB.** (A) DNA isolation and quantification workflow used for different mycobacterial strains. Total DNA yield quantified by (B) nanodrop and (C) qubit after phage D29 and CTAB extractions from identical samples (with the same number of bacterial cells). The fold increase in DNA amounts isolated by the phage-based in comparison to CTAB method are shown under the bars. Each dot represents an experimental repeat. Error bars represent the standard deviations, P-values were calculated using unpaired t-tests. ****P < 0.0001. (D) qPCR quantification of *gfp* copy numbers isolated from *Msm*::GFP and fold increase in phage D29-based DNA yields in comparison to CTAB-based yields. (E) Comparison of the extraction efficiencies of phage D29 and CTAB methods calculated relative to the theoretical genome copies in mycobacterial samples containing bacterial numbers shown on the x-axis. The data are shown as means. Error bars represent the standard deviations of two biological repeats. The P-values were calculated using unpaired t tests. *P < 0.05, **P < 0.01, ***P < 0.001.

### Supplementary Methods

#### Table S1. Key Resources

| **REAGENT or RESOURCE** | **SOURCE** | **IDENTIFIER** |
| --- | --- | --- |
| **Bacterial strains and bacteriophage (See Supplementary Table 2)** | | |
| *Mycobacterium smegmatis* mc^2^155 (WT) | (Snapper et al., 1990) | N/A |
| *Mycobacterium tuberculosis* H37Ra | ATCC | ATCC 25177 |
| *M. smegmatis* mc^2^155 pMSP12::GFP (*Msm*::GFP) | (Chan et al., 2002); provided by M. Chengalroyen | N/A |
| *M. smegmatis* mc^2^155 mScarlet (*Msm*::mScarlet) | (Kolbe et al., 2020); provided by M. Chengalroyen | N/A |
| Rifampicin-resistant *M. smegmatis* *rpoB* mutant (codon 442 A->G) | This study | N/A |
| Mycobacteriophage D29 | Hatfull Lab, University of Pittsburgh (MTA) | N/A |
| **Chemicals, dyes and reagents** | | |
| SYTOX Orange nucleic acid stain | Thermo Fisher (Invitrogen) | Cat# S11368 |
| BODIPY FL Vancomycin | Thermo Fisher | Cat# V34850 |
| Propidium iodide (PI) | Sigma-Aldrich (Merck) | Cat# P4864 |
| SYBR Gold nucleic acid stain | Thermo Fisher (Invitrogen) | Cat# S11494 |
| Middlebrook 7H9 broth base | BD Diagnostics | Cat# 271310 |
| Middlebrook 7H10 agar base | BD Diagnostics | Cat# 262710 |
| Middlebrook ADC enrichment | BD Diagnostics | Cat# 212352 |
| Glycerol | Sigma-Aldrich (Merck) | Cat# G5516-500ML |
| Tween 80 | Sigma-Aldrich (Merck) | Cat# P4780-500ML |
| Calcium chloride (CaCl_2_) | Sigma-Aldrich (Merck) | Cat# C4901 |
| Ferrous ammonium sulphate | Sigma-Aldrich (Merck) | Cat# F3754 |
| Rifampicin | Sigma-Aldrich (Merck) | Cat# R3501-25G |
| DNase I (RNase-free) | New England Biolabs | Cat# M0303 |
| HindIII restriction enzyme | New England Biolabs | Cat# R0104 |
| Power SYBR Green PCR Master Mix | Applied Biosystems | Cat# to confirm |
| 2x ddPCR Supermix for Probes (No dUTP) | Bio-Rad | Cat# 1863024 |
| Droplet Generation Oil for Probes | Bio-Rad | Cat# 1863005 |
| PrimeTime qPCR/ddPCR assay (rpoB) | Integrated DNA Technologies (IDT) | N/A |
| **Critical commercial assays / kits** | | |
| Zymo DNA Clean & Concentrator-5 | Zymo Research | Cat# D4013 |
| **Oligonucleotides** | | |
| Primers and probes (*Msm*_qGFP, *Msm*_rpoB2, Ra_23S, Ra_RD9) | This study / IDT | See **Supplementary Table S3** |
| **Software and algorithms** | | |
| Zeiss ZEN / ZEN pro | Zeiss | N/A |
| Fiji (ImageJ) | (Schindelin et al., 2012) | https://fiji.sc |
| MicrobeJ plugin | (Ducret et al., 2016) | https://microbej.com |
| FlowJo v10.8.1 | BD Biosciences | N/A |
| BD Accuri C6 software | BD Biosciences | N/A |
| QuantaSoft v1.0.596 | Bio-Rad | N/A |
| GraphPad Prism v8.0.2.263 | GraphPad Software | N/A |
| Microsoft Excel | Microsoft | N/A |
| **Others (equipment / instruments)** | | |
| CellASIC ONIX2 microfluidic system | Millipore (Merck) | N/A |
| CellASIC B04A bacteriology plates | Millipore (Merck) | Cat# B04A-03-5PK |
| Zeiss Axio Observer 7 microscope (100x 1.4 NA oil) | Zeiss | N/A |
| Axiocam 506 mono camera | Zeiss | N/A |
| BD Accuri C6 flow cytometer | BD Biosciences | N/A |
| PikoReal 96-well Real-Time PCR system | Thermo Scientific | N/A |
| QX200 Droplet Generator | Bio-Rad | N/A |
| QX200 Droplet Reader | Bio-Rad | N/A |
| T100 Thermal Cycler | Bio-Rad | N/A |
| Amicon Ultra-0.5 mL centrifugal filters (Ultracel-30K) | Merck Millipore | Cat# UFC5030 |
| 0.22 µm syringe / membrane filters | Millipore (Merck) | Cat# SLGV033RS |
| Micro-emulsifying needle (25G, 4-inch) | Sigma-Aldrich | Cat# CAD7974 |

#### Table S2. Mycobacterial strains used in this study

| **Bacterial strain** | **Description** | **Assay** | **Reference** |
| --- | --- | --- | --- |
| *Mtb* H37Ra | Virulence-attenuated (Biosafe) strain of *Mtb* ATCC 25177. | Used in measuring phage lysis and DNA extraction in *Mtb* | ATCC |
| *Msm* mc^2^155 | WT *Msm* strain. | Used for phage amplification, DMN-Tre labelling and DNA extraction efficiency investigations | (Snapper et al., 1990) |
| *Msm*::GFP | mc^2^155 derivative harbouring pMSP12::GFP vector integrated in the genome. | Used to investigate single-cell phage adsorption, lysis and DNA extraction efficiency | (Chan et al., 2002) |
| *Msm*::mScarlet | mc^2^155 derivative containing genome-integrated mScarlet fluorescent protein. | Used to investigate mycobacterial permeation and uptake of BODIPY FL Van | (Kolbe et al., 2020) |
| *Msm rpoB* | mc^2^155 derivative carrying a *rpoB* mutation (codon 442 A->G) conferring rifampicin resistance. The mutant was isolated from rifampicin resistant mc^2^155 colonies and confirmed by Sanger sequencing. | Used to demonstrate phage D29 plaque formation/lysis on rifampicin-resistant mycobacteria | (This study) |

#### Table S3. Primers and probes used for qPCR and ddPCR quantification

| **Oligo ID** | **5’ → 3’ sequence** | **Assay** | **Description** | **Reference** |
| --- | --- | --- | --- | --- |
| *Msm*_qGFP 200_Fwd | CCCAGATCACATGAAACGGCA | qPCR | Relative gDNA quantification in *Msm*::GFP targeting *gfp* gene | (This study) |
| *Msm*_qGFP 500_Rvs | ACGTGTCTTGTAGTTCCCGTC | qPCR | Relative gDNA quantification in *Msm*::GFP targeting *gfp* gene | (This study) |
| *Msm*_rpoB2_916_Fwd | AAGGTCAACAAGAAGCTGG | ddPCR | Absolute gDNA quantification in *Msm* targeting 93bp of *rpoB* gene | (This study) |
| *Msm*_rpoB2_992_Rvs | CAGGTACTCGATGGTCG | ddPCR | Absolute gDNA quantification in *Msm* targeting 93bp of *rpoB* gene | (This study) |
| *Msm*_rpoB2_957_probe | CACCAGCTCGACGCTGACCG | ddPCR | PrimeTime™ HEX-based fluorophore reporter for absolute quantification of gDNA in *Msm* targeting *rpoB* gene | (This study) |
| Ra_23S_Fwd | GCAGCGAAAGCGAGTCTGA | ddPCR | Absolute gDNA quantification in H37Ra targeting 23S rRNA gene in gDNA | (Walter et al., 2021) |
| Ra_23S_Rvs | CCAGAACACGCCACTATTCACA | ddPCR | Absolute gDNA quantification in H37Ra targeting 23S rRNA gene in gDNA | (Walter et al., 2021) |
| Ra_23S_Probe | AGGGCGACCCACACGCGC | ddPCR | HEX-based fluorophore reporter for absolute quantification of gDNA in H37Ra targeting 23S rRNA gene | (Walter et al., 2021) |
| Ra_RD9_Fwd | TGAGTGGCGATGGTCAACAC | ddPCR | Absolute gDNA quantification in H37Ra targeting RD9 region in genomic DNA (gDNA) | (Wood et al., 2016) |
| Ra_RD9_Rvs | GATGGCGTTCGGAAAGAAAC | ddPCR | Absolute gDNA quantification in H37Ra targeting RD9 region in gDNA | (Wood et al., 2016) |
| Ra_RD9_Probe | ACTACGCGGCTTAGTG | ddPCR | FAM-based fluorophore reporter for absolute quantification of gDNA in H37Ra targeting RD9 region | (Bunyasi et al., 2022) |

### Additional methods

#### Determination of MOI

Determination of optimal MOI is critical for effective bacterial infection and in vitro phage lysis. A high MOI could increase the probability for the development of bacterial resistance, whereas a low MOI may not produce adequate phage titer to trigger infection (Międzybrodzki et al., 2023). For these reasons, the dose of phage required for optimal bacterial lysis in was investigated using *Msm*.

The WT *Msm* cells from glycerol stocks were cultured to exponential phase (OD_600_ = 0.3-04) and passaged twice. Aliquots of *Msm* (1 mL) culture were infected with 100 µL of phage particles at an estimated MOI 1, 10, 100 and 1000. The infected bacteria were incubated for 12 hr at 37°C in a shaking incubator at 150 rpm and then spread on 7H10 plates. In parallel, an equivalent volume of uninfected *Msm* culture was spread on 7H10 control plates. The plates were incubated for 5 days and the CFUs calculated. The CFU from phage infected cultures were assessed relative to the CFUs from control plates to determine survival.

#### ddPCR cycling conditions and reaction compositions

ddPCR reactions were used to quantify the gDNA isolated by CTAB and phage D29 methods in (i) *Msm* by targeting the *rpoB* gene and in (ii) *Mtb* (H37Ra) by targeting 23S ribosomal RNA (rRNA) gene and Region of Difference 9 (RD9). Freshly isolated gDNA originating from samples with known number of bacterial cells (320, 3200, and 32000 in *Msm*) and (175, 17500, and 175000 in *Mtb* H37Ra), as determined by flow cytometry were used as ddPCR templates. For *Msm* ddPCR, 1xÍreactions of 22 µL total volume contained 1 µL of *Hin*dIII restriction enzyme, 11 µL of 2xddPCR SuperMix (No dUTP), 2.2 µL of 10xPrimeTime Assay containing *Msm*_rpoB2 primers and probe from IDT (**Table S3**), 2 µL of template and 5.8 µL of nuclease-free water. For *Mtb* ddPCR, 1xreactions of 22 µL total volume contained 1 µL of *Hin*dIII restriction enzyme, 11 µL of 2xddPCR SuperMix (No dUTP), 0.6 µL of 30 µM forward primers, 0.6 µL of 30 µM reverse primers, 0.6 µL of 10 µM probes (**Table** **S2**), 2 µL of template DNA, and 6.8 µL of nuclease-free water. The reactions were emulsified with droplet generator oil (Bio-Rad) and partitioned into droplets using a QX200 Droplet Generator (Bio-Rad). Thermal cycling was performed in a T100 Thermal Cycler (Bio-Rad), thermal cycling conditions were as follows: 1 cycle at 95°C for 10 min, 40 cycles at 94°C for 30 s, 58.3°C for 60 s, and 98°C for 10 min. The reactions were held at 4°C overnight after thermal cycling to stabilize the droplets. The droplets were then read in a QX200 Droplet Reader (Bio-Rad), and data exported to QuantaSoft Software version for analysis. Further analyses were performed in Excel and GraphPad Prism.

#### Quantification of *Msm*::GFP DNA by qPCR

DNA recovered from defined numbers of GFP expressing *Msm* cells, was quantified by qPCR using primers targeting the chromosomally integrated *gfp* gene (**Table S3**). Reactions were prepared using the Power SYBR® Green PCR Master Mix and quantification performed against a serially diluted gDNA standard. Cycling conditions and reaction compositions are provided in detail below. To examine the functionality of the qPCR assay, a 10-fold serial dilution series of an 80 ng/µL gDNA stock was performed to 10^-6^ (8 ng/µL, 0.8 ng/µL, 0.08 ng/µL, 0.008 ng/µL, 0.0008 ng/µL and 0.00008 ng/µL) , and the samples loaded on the assay for the assessment of primer binding and the cycle threshold (CT) values. The assays were prepared in 20 µL reactions that consisted of the following: 10 µL SYBR® Green PCR master mix, 200 nM of the *Msm*_qGFP primers, 2 µL of template DNA and nuclease-free water to make the reactions to 20 µL. qPCR reactions were carried out on the PikoReal® 96-well Real-Time PCR system (Thermo Scientific) with the cycling conditions set as follows: initial denaturation at 95°C for 10 min, 40 cycles of denaturation at 95°C for 30 s, annealing and extension at 60°C for 1 min, followed by data acquisition. Data were analysed in Excel and GraphPad Prism.

#### Live-cell time lapse microscopy

Live-cell imaging was performed using a CellASIC ONIX2 microfluidic system with B04A bacteriology plates mounted on the Zeiss Axio Observer 7 system. The experimental protocols are described in detail below.

Mycobacterial cells were grown to OD_600_ = 0.3-0.4, filtered through 5 µm filters to create single cell suspensions and remove any debris, and loaded on the CellASIC plates. Experimental treatments were loaded based on protocols used as detailed below. Plates were then sealed and mounted on the microscope stage. Bacterial loading cycles were performed using the pressure-driven protocol as specified by the manufacturer. On the microscope, an imaging session was selected for each experiment, and the imaging channels and exposure settings selected. The ‘tiles setup’ menu was selected as the focus method with software Autofocus as an additional action. Autofocus software was set in the range of 40 µm to permit the software to search 20 µm above and below the specified Z-plane to bring samples into focus. Time-lapse settings were selected followed by selection and addition of the regions of interest for imaging in live mode under 100X oil immersion objective. The Z-positions for all the selected tiles were verified and manually adjusted where necessary to ensure that the images were in focus. The CellASIC software and timelapse experiment were started simultaneously. Temperature was maintained at 37°C throughout the experiments. Images were captured using Zeiss ZEN pro software (Zeiss) and exported to either Fiji software or MicrobeJ plugin for analysis.

Specific live cell time lapse experiments were performed as follows:

i. **GFP release experiment**: *Msm*::GFP cells were loaded in cycles of 30 s until ≥5 bacteria were trapped in the imaging chambers. The experimental treatments in columns 1, 2, and 3 of the plate were perfused for durations of 1 hr (phage medium only), 6 hr (phage), and 17 hr (phage medium only) respectively (**Figure SM1A**). GFP signal was recorded using the green fluorescent protein detector (excitation wavelength 450-490 nm and emission wavelength 500-550 nm) and filtered at 450-488 nm. An exposure time of 150 ms was maintained throughout the experiment. For each experimental condition five fields of view were captured every 30 min.
ii. **GFP release and uptake of propidium iodide (PI):** *Msm*::GFP bacteria were loaded in cycles of 30 s until the desired number of bacteria was trapped in the imaging chambers. Experimental and control conditions on the plate were perfused for 1 hr (phage medium only), 1 hr (phage), 1.5 hr (phage medium only), 0.5 hr PI and 1.5 hr (phage medium only), respectively (**Figure SM1B**). GFP excitation, filtering and exposures were as above. PI fluorescence was excited using Alexa Fluorescence 568 detector (excitation wavelength 550-580 nm and emission wavelength 590-650 nm) and filtered at 540-570 nm, exposure time was maintained at 150 ms throughout the experiment. For each experimental condition five fields of view were captured every 6 min.
iii. **Uptake of BODIPY FL Vancomycin (BODIPY-FL-van):** *Msm*::mScarlet bacteria (Kolbe et al., 2020) were loaded in the imaging chambers in cycles of 30 s until the desired number of cells was trapped in the imaging chambers. The experimental and control treatments on the plate were perfused for 1 hr (phage medium only), 1 hr (phage), 4 hr (phage medium with BODIPY-FL-van), and 4 hr (phage medium only), respectively (**Figure SM1C**). Red fluorescence (mScarlet) was excited using Alexa Fluorescence 568 detector (excitation wavelength 550-580 nm and emission wavelength 590-650 nm) and filtered at 540-570 nm, exposure time was maintained at 200 ms throughout the experiment. Green fluorescence (BODIPY-FL-van) was excited using a Green Fluorescent Protein detector (excitation wavelength 450-490 nm and emission wavelength 500-550 nm) and filtered at 450-488 nm, the exposure time was maintained at 150 ms throughout the experiment. For each experimental condition five fields of view were captured every 30 min.

#### Supplementary Methods Figures

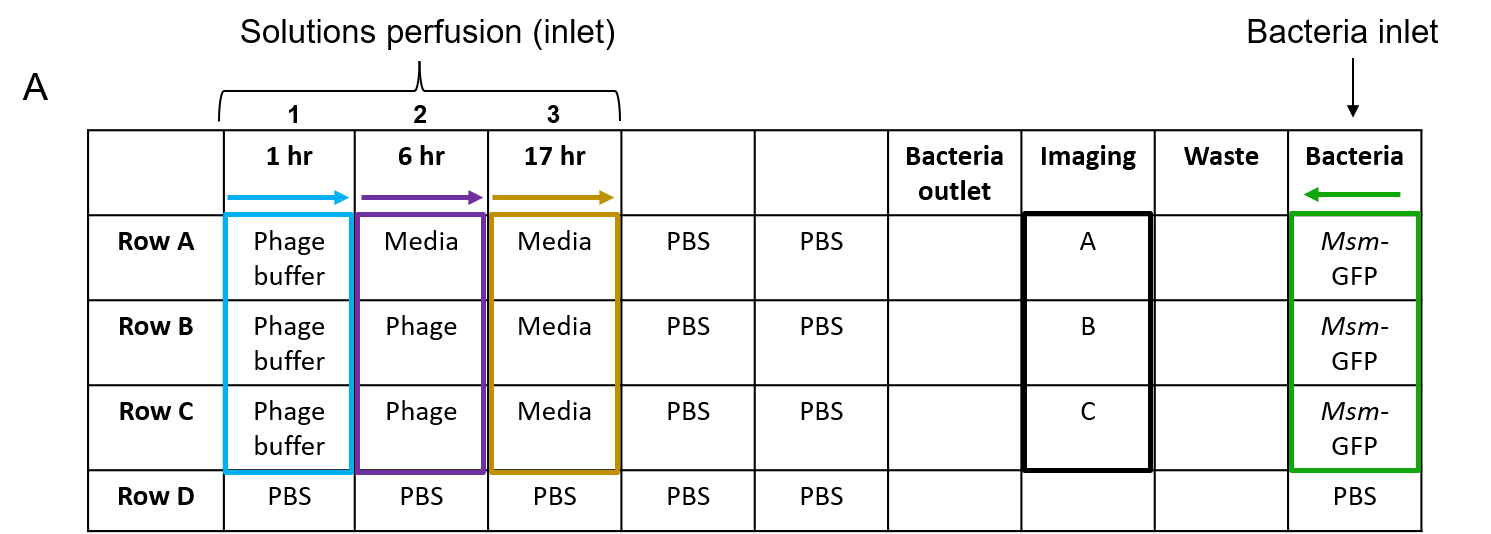

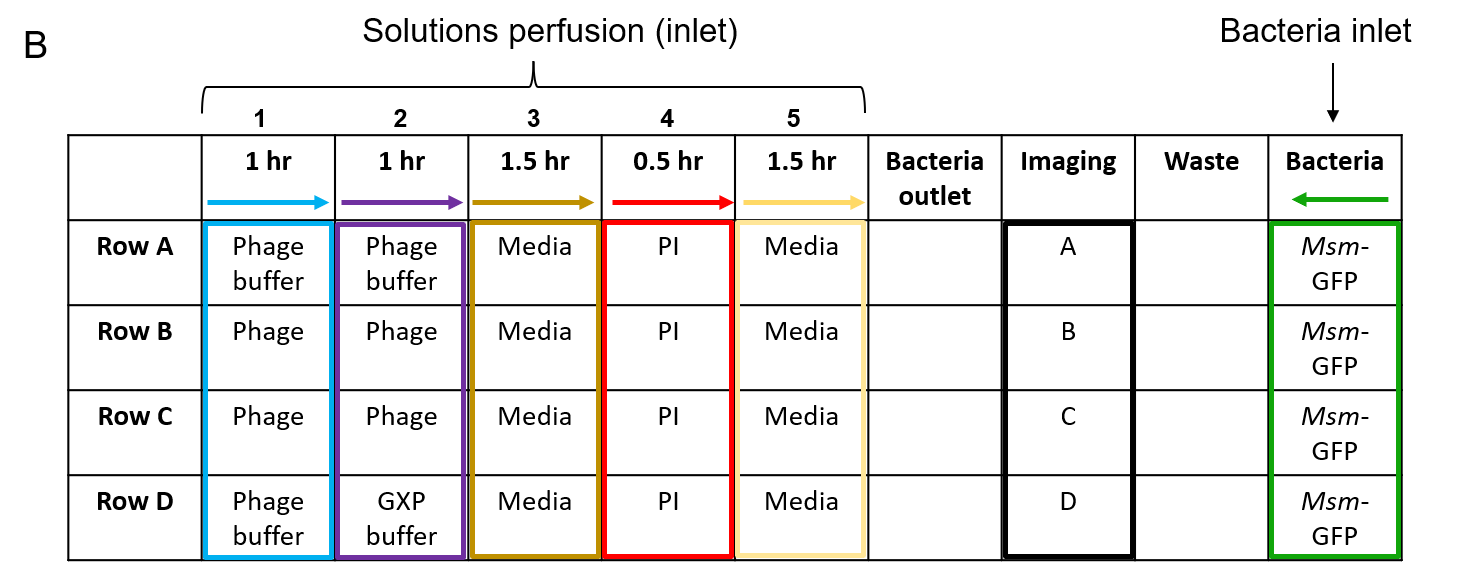

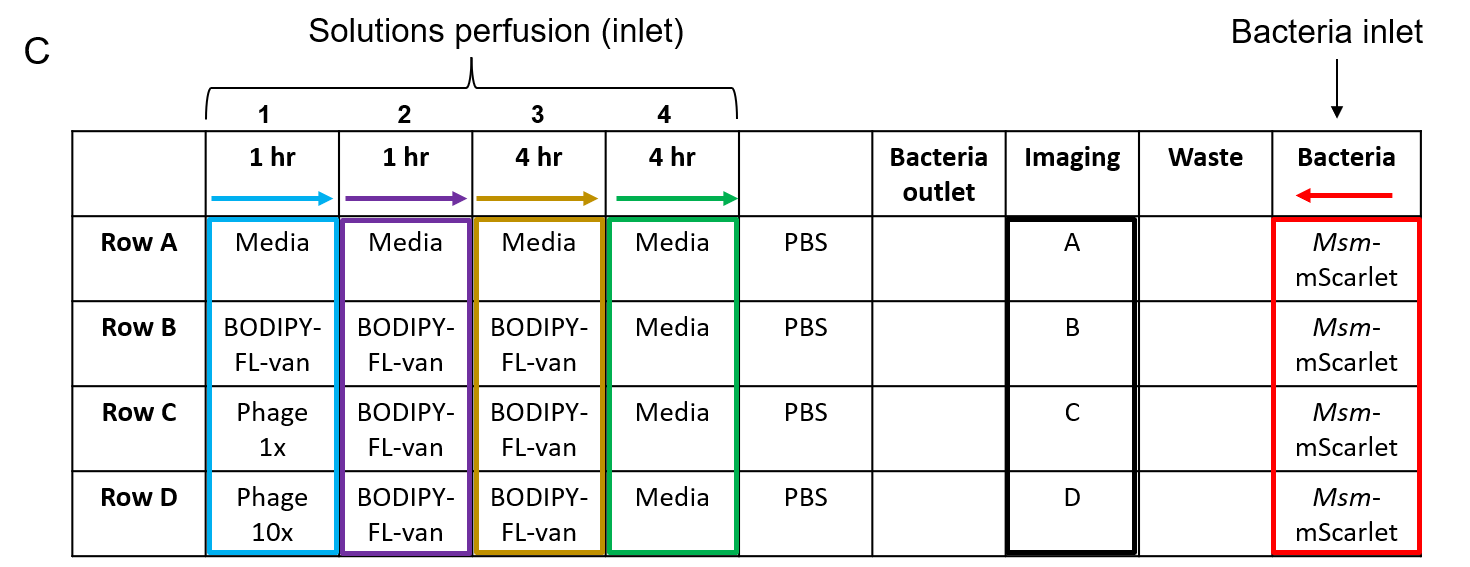

**Figure SM1: CellASIC bacterial plate maps and experimental conditions applied for different experiments.** (A) Imaging GFP release. Perfusion of phage buffer only for 1 hr (blue), perfusion of phage buffer and phage (purple) for 6 hr and, lastly, 17 hr of medium perfusion (brown). (B) Imaging GFP release and uptake of PI. Perfusion of phage and phage buffer for 1 hr (blue), perfusion of phage buffer, phage and GXP buffer for 1 hr (purple), perfusion of media for 1.5 hr (brown), perfusion of PI for 0.5 hr (red) and 1.5 hr of media perfusion (yellow). (C) Imaging the uptake of BODIPY-FL-van. Media, BODIPY-FL-van, 1X phage and 10X phage solutions were perfused for 1 hr (blue), media and BODIPY-FL-van were perfused for 1 hr (purple) followed by an extra 4 hr perfusion (brown), and lastly, media was perfused for 4 hr (green).
